# Molecular Determinants of Receptor-Specific Membrane Interactions in EphA1 and EphA2 from Coarse-Grained Simulations

**DOI:** 10.64898/2025.12.23.696317

**Authors:** Amita R. Sahoo, Nisha Bhattarai, Matthias Buck

## Abstract

Ephrin receptors (Ephs) are receptor tyrosine kinases that regulate cellular growth, differentiation, and motility. EphA2, often overexpressed in cancer, is notable for its ligand-independent activation, which drives pro-oncogenic signaling distinct from the canonical, ligand-dependent pathway that restricts cell movement. While ligand binding induces extracellular clustering, kinase activation depends on dimerization within the transmembrane (TM) region. EphA1 and EphA2 differ substantially in their function, likely due to differences in their TM, but more-so in their juxtamembrane (JM), and membrane-proximal fibronectin type III (FN1/FN2) domains. How these latter two regions modulate dimerization has not yet been investigated. To address this, we performed extensive coarse-grained (CG) simulations using Martini 3 in an anionic POPC/PS/PIP₂ model of the plasma membrane. Both receptors formed stable TM dimers, though EphA1 favored a symmetric AXXXGXXXG-centered interface, whereas EphA2 favoured an alternate leucine zipper interface. All protein constructs sampled multiple configurations, reflecting substantial intrinsic variability. AlphaFold3 does not yield reliable predictions and cannot account of effects of the lipid bilayer. In the CG simulations, basic residues in the JM region remained membrane-bound, and the EphA2 FN domain displayed sustained PIP₂ interactions, consistent with previous observations. Notably, constructs with the FN2 domain alone restricted TM association in both receptors, whereas inclusion of the second FN1 domain restored dimerization but produced receptor-specific extracellular interfaces. These differences arise from distinct FN1-FN2 linker flexibilities, a different level sequence conservation of EphA1 compared to several other Ephs and FN-domain membrane contacts, which together shape TM geometry and lipid engagement. Our results predict how TM and TM-proximal elements cooperatively tune dimerization in Eph receptors. This work offers a molecular rationale into the divergent activation behaviors of EphA1 and EphA2 and provides testable models relevant to cancer biology and Eph-driven signaling.

## Introduction

Cancer remains a major global health challenge and a leading cause of death worldwide.^1,2^ It arises from the dysregulation of fundamental cellular processes such as the cell cycle, cellular membrane lipid composition, apoptosis, and cell migration processes that are normally governed by intricate, interconnected signaling pathways.^3,4^ A prominent class of signaling proteins involved in these pathways are receptor tyrosine kinases (RTKs), which often act as proto-oncogenes and contribute to carcinogenesis by amplifying signaling activity.^5,6^ Among the RTKs, Eph receptors represent the largest subgroup and certain members have garnered significant attention for their roles in tumorigenesis.^7–9^ These receptors, through their interaction with membrane-bound ephrin ligands, regulate diverse cellular behaviors, including cell-cell communication and migration. Expression levels of Eph receptors vary with cancer type and stage, suggesting their potential as diagnostic or prognostic markers. For instance, reduced EphA1 expression is associated with advanced stages of colorectal and non-melanoma skin cancers, while elevated levels are observed in locally invasive colorectal tumors.^10,11^ Similarly, EphA2 expression fluctuates across different cancer stages and the receptor is frequently overexpressed in numerous cancer types and cancer-derived cell lines. Given its widespread expression and upregulation in tumors, EphA2 has emerged as a promising biomarker for clinical cancer management. Furthermore, the differential expression patterns of EphA1 and EphA2 in normal versus cancerous tissues underscore their relevance as therapeutic targets.^12–15^ Despite their highly conserved domain organization and substantial sequence similarity, EphA1 and EphA2 exhibit distinct biological functions and signaling characteristics. EphA2 is well recognized for its ligand-independent signaling and oncogenic activity in numerous cancers, whereas EphA1 has generally been associated with different physiological and pathological roles, including the regulation of cell adhesion and migration. The structural determinants responsible for these receptor-specific functional differences remain poorly understood, motivating comparative studies of their membrane organization and dimerization mechanisms.

Eph receptor activation is typically initiated through interaction with membrane-anchored ephrin ligands. Ligand binding induces receptor dimerization, which several reports suggest is followed by further oligomerization into higher-order clusters.^16,17^ Within these clusters, Eph receptors and associated downstream signaling proteins undergo tyrosine phosphorylation, either through receptor autophosphorylation or via recruitment of SH2 domain-containing effectors to phosphotyrosine sites on Eph receptors.^18^ Several studies have proposed that EphA2 receptors can undergo ligand-independent dimerization when their local concentration in the plasma membrane becomes sufficiently high. However, a point mutation of P460L in the EphA1 ectodomain is also known to promote clustering and activation of EphA1 receptors independently of ligand binding.^19–21^ In addition to ligand-mediated activation, proteolytic cleavage of Eph receptors plays an important role in regulating their function. EphA2, as well as the related receptors EphA4 and EphB2, undergo extracellular cleavage by MT1-MMP between their fibronectin type III domains (FN1/FN2), and by the TF/FVIIa complex within the ligand-binding domain^22–25^ with possible relevance to neurodegeneration and the extracellular matrix transition (EMT) in cancers where similar fragments are generated. Notably, the EphA2 constructs analyzed in our study mimic the size and domain composition of these physiologically generated cleavage fragments. This relevance provides a mechanistic rationale for our construct design for truncated extracellular regions coupled to TM and JM dimerization. While numerous structural studies have characterized individual domains of receptor tyrosine kinases including the ectodomain, transmembrane (TM) domain, and kinase domain the integration of the behaviour of these elements into fuller-length, membrane-embedded model of the receptor remains a major challenge.^26^ AlphaFold3, a machine learning based protein structure prediction algorithm has revolutionized structural biology as it reliably predicts structures of well folding protein domains and protein-protein complexes.^27^ However, it does fail for certain systems: while it gave accurate predictions for most TM region dimers,^28^ surprisingly it is unable to predict reasonable structures for EGFR and EphA2 kinase domain dimers^29^ and as we show in this report it also does not predict reasonable structure for EphA1 and A2 protein fragments where soluble domains, FN and JM regions surround the transmembrane segment. By contrast, molecular dynamics (MD) simulations have provided a powerful framework to investigate the structural dynamics of transmembrane proteins and their interactions.^30–32^Previous studies have investigated the homodimerization of their TM domains of EphA1 and EphA2 ^28,33^ as well as the interactions of the membrane-proximal FNIII domain of EphA2 (FN2) with the lipid bilayer,^28,34^ providing important insights into the configurational flexibility, membrane association and lipid-mediated regulation of these isolated domains. More broadly, both computational studies of receptor tyrosine kinases (RTKs)^30,31,35^and experimental investigations^36^ have demonstrated that RTK ectodomains can directly interact with lipid bilayers. However, despite these advances, the membrane interactions and conformational dynamics of larger Eph receptor constructs containing both extracellular fibronectin domains together with the transmembrane and juxtamembrane regions remain largely unexplored.

In this study, we investigated the ligand-independent dimerization of the EphA1 and EphA2 receptors, with a focus on their transmembrane (TM) regions when they are joined to their membrane-proximal domains (Fig. S1A). To dissect the role of different structural elements, we modeled three receptor constructs: the TM alone, FN2-TM-JM, and FN1-FN2-TM-JM and embedded them in a physiologically approximate, mixed anionic lipid bilayer, consisting of 80% POPC, 15% POPS and 5% PIP2 (Fig. S1B and C). Coarse-grained molecular dynamics simulations using the Martini3 force field allowed us to compare the intrinsic dimerization propensities of EphA1 and EphA2 and to evaluate how extracellular domain orientation, linker flexibility, and lipid interactions influence TM association. Finally, we discuss how this stepwise approach highlights sequence- and domain-specific determinants governing receptor organization, extracellular domain flexibility, membrane interactions, and transmembrane dimerization. These structural differences provide mechanistic insights into how EphA1 and EphA2 may exhibit distinct functional properties.

## Results

### TM-only dimerization of EphA1 & EphA2

To investigate the intrinsic dimerization propensities of the EphA1 and EphA2 transmembrane (TM) domains, we modeled three receptor fragments of varying lengths (Fig. S1B). Beginning with the smallest construct comprising only the TM domain, coarse-grained (CG) molecular dynamics simulations were conducted using the Martini3 force field with an anionic-enriched mixed lipid bilayer containing POPC (80%), POPS (15%), and PIP2 (5%). This composition has been used before as an approximate model for the plasma membrane of eukaryotic cells and inclusion of anionic lipids, particularly PIP2, reflects on the importance of lipid-protein interactions in modulating receptor function. We used a bilayer with a symmetric composition of lipids, as loss of asymmetry is known to occur in many cancers.^37,38^ In each system, two identical TM helices were embedded in the bilayer in parallel orientation with respect to the membrane normal and initially separated by 5.0 nm to prevent bias from starting configurations (Fig. S1C). Each system was simulated in quadruplicate for 4 μs per replicate, totalling 16 μs of simulation per construct. During the simulations, spontaneous association of the TM helices was typically observed within 200 to 1000 ns, after which the dimers remained stably associated. The time evolution of the center-of-mass distance between TM helices showed a clear convergence for both EphA1 and EphA2 systems (Fig. S2A & B), supporting the formation of stable dimeric states. To systematically assess the dimer configurations, all replicates were concatenated and projected onto 2D plots showing population of the inter-helical crossing angle (θ) versus inter-helical center-of-mass distance (d) (Fig. S2C & D). The EphA1 dimer population was primarily clustered within distance of 0.6 to 0.9 nm and crossing angle of –75° to +90°, while the EphA2 dimers exhibited a broader distance range, spanning 0.6 to 1.1 nm and but lesser crossing angles of –40° to +75°. This spread in configurational space for both EphA1 and EphA2 suggests differences in structural preferences, a likely polymorphism in their dimer interface (see below).

We performed hierarchical clustering using a 0.6 nm backbone RMSD cutoff to identify the most representative dimer configurations (Fig. S3). Analysis of the top clusters per receptor revealed distinct interaction interfaces. Therefore, an average residue-residue contact maps were computed for the dimeric ensembles of each receptor (Fig. S2E & F), revealing distinct interaction patterns. For EphA1, the transmembrane dimers mainly formed symmetric arrangements centered on the conserved AXXXGXXXG motif. The glycine residues in this motif allow the helices to pack closely, producing interfaces that closely match those seen in the NMR structure^39^ (Fig. 1A). In contrast, EphA2 transmembrane dimers showed greater structural variability. Their interfaces involved both the N-terminal GXXXG motif and a second region resembling a leucine-zipper like hydrophobic motif. This suggests that EphA2 dimerization may rely on a combination of glycine-mediated helix packing and additional hydrophobic side-chain interactions, consistent with features observed in the NMR structure^40^ (Fig. 1B). The importance of these motifs in EphA2 have been explored experimentally through several mutagenesis studies.^41,42^ We also observed dynamic interconversion between left-handed and right-handed dimer configurations (more evident in EphA1 compared to EphA2) (Fig. S2C-D and S4), which reflect configurational preferences in the TM helix interfaces and support the idea of functionally relevant dimer polymorphism. Such polymorphisms may be essential for accommodating different signaling states or responding to distinct lipid microenvironments. These observations are consistent with our previous CG Martini 3 studies of Eph TM domain dynamics in DMPC bilayers, confirming that both EphA1 and EphA2 TM domains form stable dimers, but with distinct interfacial motifs and configurational landscapes.^28^ Importantly, the preference for different dimer interfaces between EphA1 and EphA2 and ultimately their differences in their amino acid sequences underlie their different signaling behavior and sensitivity to membrane context.

**Figure 1.**
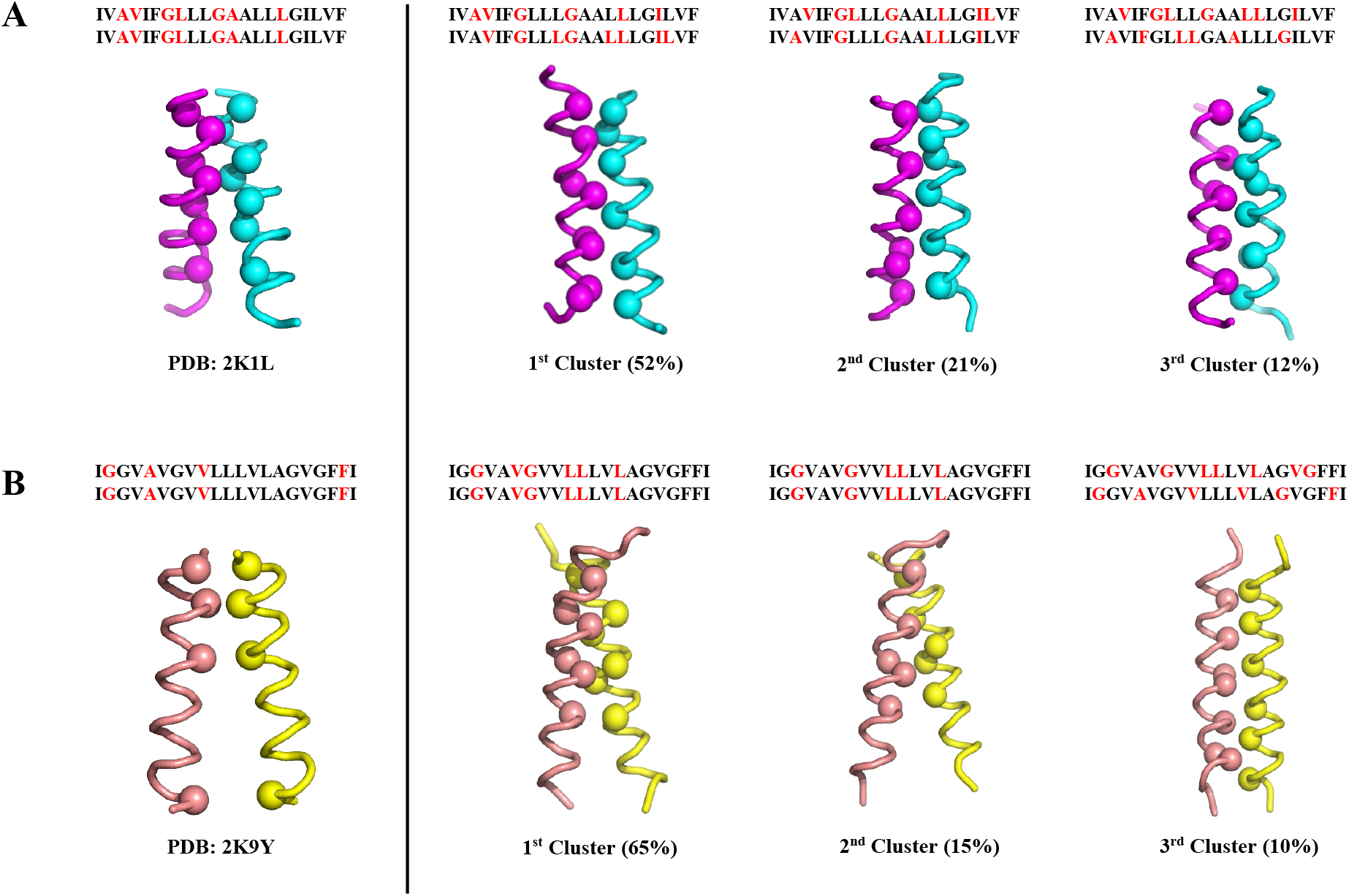
Comparison of most populated clusters of the TM-only construct dimer of EphA1 (A) and EphA2 (B) with the corresponding NMR structures. All the conformations are shown in ribbon representation and the Cα atoms of the TM interface interacting residues are shown as spheres. Contact residues are highlighted as red. Population of individual clusters of the TM dimers of both EphA1 and EphA2 are shown here. Clustering of both EphA1 and EphA2 dimers are done with 6Å cutoff.

### FN2-JM modulation of EphA TM dimerization

To examine how membrane-adjacent extracellular and intracellular elements influence TM dimerization, we extended our CG models to include the second, most membrane proximal fibronectin type III (FN2) domain at the N-terminus and the full juxtamembrane (JM) domain at the C-terminus of the TM segment for both EphA1 and EphA2 (Fig. S1B). These domains are anticipated to modulate the orientation and association of TM helices by imposing longer range spatial and/or close-contact steric constraints. The simulations were performed, as for the TM region above, but as eight independent replicas of 4 µs each. In contrast to the TM-only systems, the inter-helical distances remained significantly larger across all trajectories when the FN2 and JM domains were present (Fig. S5), indicating that the TM helices do not stably dimerize in these constructs with an extracellular FN2 domain. This demonstrates that the adjacent domains sterically hinder TM-TM interactions. To further explore this, we performed clustering of all simulation frames using a 1.0 nm backbone RMSD cutoff. Representative structures from the most populated clusters for EphA1 and EphA2 are shown in Fig. 2A-B.

**Figure 2.**
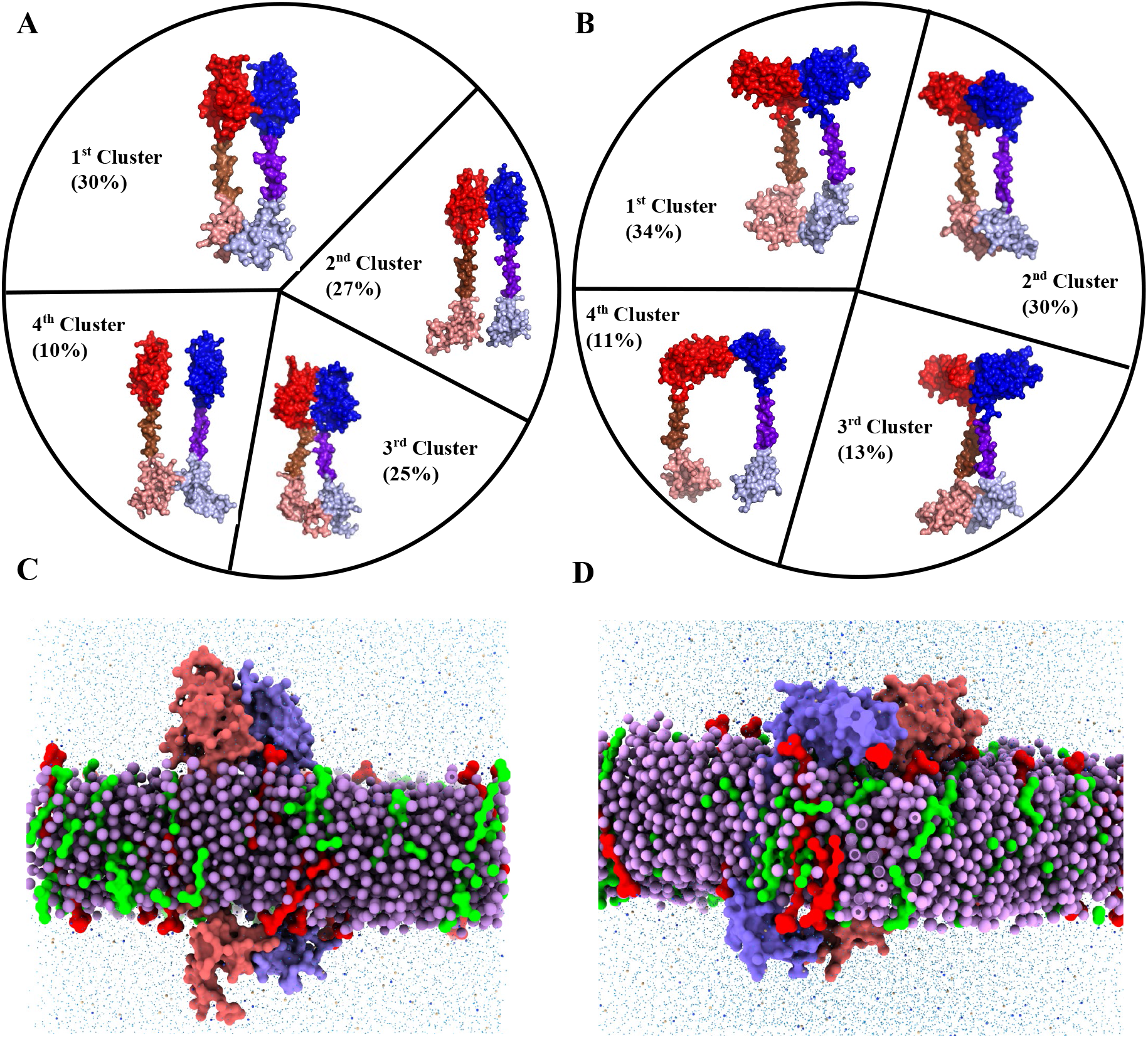
Comparison of the most populated clusters of the FN2-TM-JM construct dimers of EphA1 (A) and EphA2 (B). Structures are shown in surface representation. Monomers are designated as chain A and chain B throughout. In chain A, the FN2, TM, and JM regions are colored red, chocolate, and salmon, respectively; in chain B, they are colored blue, purple-blue, and light blue. Clustering was performed using a 1 nm cutoff, based on the final 2.5 µs of each trajectory from eight simulations. (C, D) Representative structures of the dominant clusters of EphA1 (C) and EphA2 (D) dimers in a mixed membrane. Chain A and chain B are shown as orange-red and purple surfaces, respectively. PIP2, POPS, and POPC are shown as red, green, and purple spheres.

For EphA1, the FN2 domains adopt an upright configuration, projecting away from the membrane and exhibiting minimal lipid contact (Fig. 2A). This arrangement is evident in the membrane snapshot of the EphA1 dimer (Fig. 2C). Analysis of FN2-lipid contacts revealed several residues, including R454, K457, K458, R461, R534, T535, S536, P537, and P538, engaging with PIP2 lipids (Fig. S6A). The extended, membrane-distal positioning of FN2 appears to spatially separate and reorient the TM segments, thereby preventing persistent TM– TM engagement throughout the simulations.

EphA2 displays markedly different behavior. Its FN2 domains frequently lie adjacent to or along the membrane surface, likely stabilized by electrostatic interactions with anionic lipids such as PIP2 or POPS (Figs. 2B and 2D). This membrane-proximal orientation of FN2 may restrict the lateral mobility of the TM helices or impose a tilt that impairs productive dimer formation. Interestingly, in approximately 13% of the sampled conformations, specifically in the third most populated cluster, the TM helices do form a dimer interface, suggesting a partial or transient engagement despite the presence of FN2 and JM domains. In this context, several residues of the FN2 region including N435, K441, R443, R447, R465, T501, T502, K520, G531, S532, and G533 were observed to interact with PIP2 (Fig. S6B). Notably, some of these residues (e.g., K441, R443) were previously identified by Chavent et al. (2016) as key mediators of EphA2-PIP2 interactions^34^ further validating our observations. Together, these results highlight a key structural difference between EphA1 and EphA2 in their membrane-proximal architectures. The distinct membrane associations of the FN2 domain and likely the JM region modulate the spacing and orientation of the TM helices and thereby alter their ability to form stable dimers.

### Restoration of TM dimerization by incorporation of the second FN, the FN1 domain

To evaluate how further extension of the extracellular region influences transmembrane dimerization, we added (counting from the protein’s N-terminus) the first fibronectin type III domain (FN1) to the FN2-TM-JM constructs of both EphA1 and EphA2 (Fig. S1B). As above eight independent 4-µs replicas were performed for each receptor. Analysis of the interhelical distances between TM regions (Fig. S7A, B) revealed a notable increase in TM dimerization compared to the FN2-TM-JM systems. TM-TM interactions were observed in 60% of simulations (5 out of 8) for EphA1 and in 87% (7 out of 8) for EphA2, indicating that the presence of the second FN1 domain can, in many cases, restore or even promote TM association despite the additional extracellular mass. The 2D plots (Fig. S7C, D) showed TM dimeric distributions similar to those found in the TM-only constructs (Fig. S2C, D), suggesting that the second FN domain does not sterically block TM packing to the same extent as in the FN2-TM-JM construct. Additionally, we compared the positions of the transmembrane (TM) interface residues across all three constructs for both EphA1 and EphA2 (Fig. S8). In the presence of only the FN2 domain, the orientation of FN2 appears to alter the positioning of the TM helices, rotating the interface residues outward and thereby preventing stable TM dimerization, whereas inclusion of both FN1 and FN2 restores a favorable helical orientation consistent with productive TM dimerization.

Clustering of dimeric frames using a 2.0 nm backbone RMSD cutoff (Fig. 3A-B) revealed that addition of the FN1 domain significantly alters extracellular domain geometry and rigidity. In EphA1, FN1 and FN2 adopt a relatively rigid, membrane adjacent configuration, lying flat along the bilayer rather than projecting upward as in the FN2-TM-JM construct. This configuration enables extensive lipid interactions for both FN domains. In both receptors, FN1/FN2 contacts frequently contribute directly to the dimer interface, displaying diverse interaction modes including side-side, face-face, and opposite-facing arrangements. Notably, interactions of the respective TM and JM domains co-occur with FN-domain contacts in a large fraction of dimers ∼72% for EphA1 and ∼90% for EphA2 showing that extracellular domain engagement is not mutually exclusive with TM and JM association in these extended constructs. Lipid-interaction patterns were receptor-specific, several residues within both FN1 and FN2 engage PIP2 in both receptors (Fig. 3C-D). Notably, EphA1 exhibits substantially higher FN1-PIP2 interaction frequency compared with EphA2 (Fig. S9A-B), reflecting its broader membrane-apposed FN-domain surface. This differential behavior is also evident in the surrounding lipid environment: PIP2 molecules show pronounced local enrichment and clustering around EphA1 relative to EphA2 (Fig. S9C). Enhanced PIP2 recruitment likely stabilizes the membrane-proximal configuration of EphA1’s FN domains and may contribute to the more symmetric and extensive extracellular dimer interfaces observed for this receptor. Moreover, comparison of JM-PIP2 contacts reveals that FN1-FN2 domain dimers engage PIP2 more extensively than single FN2 dimers in both EphA1 and EphA2 (Fig. S10), suggesting that both extracellular and intracellular domains contribute synergistically to TM dimer stabilization.

**Figure 3.**
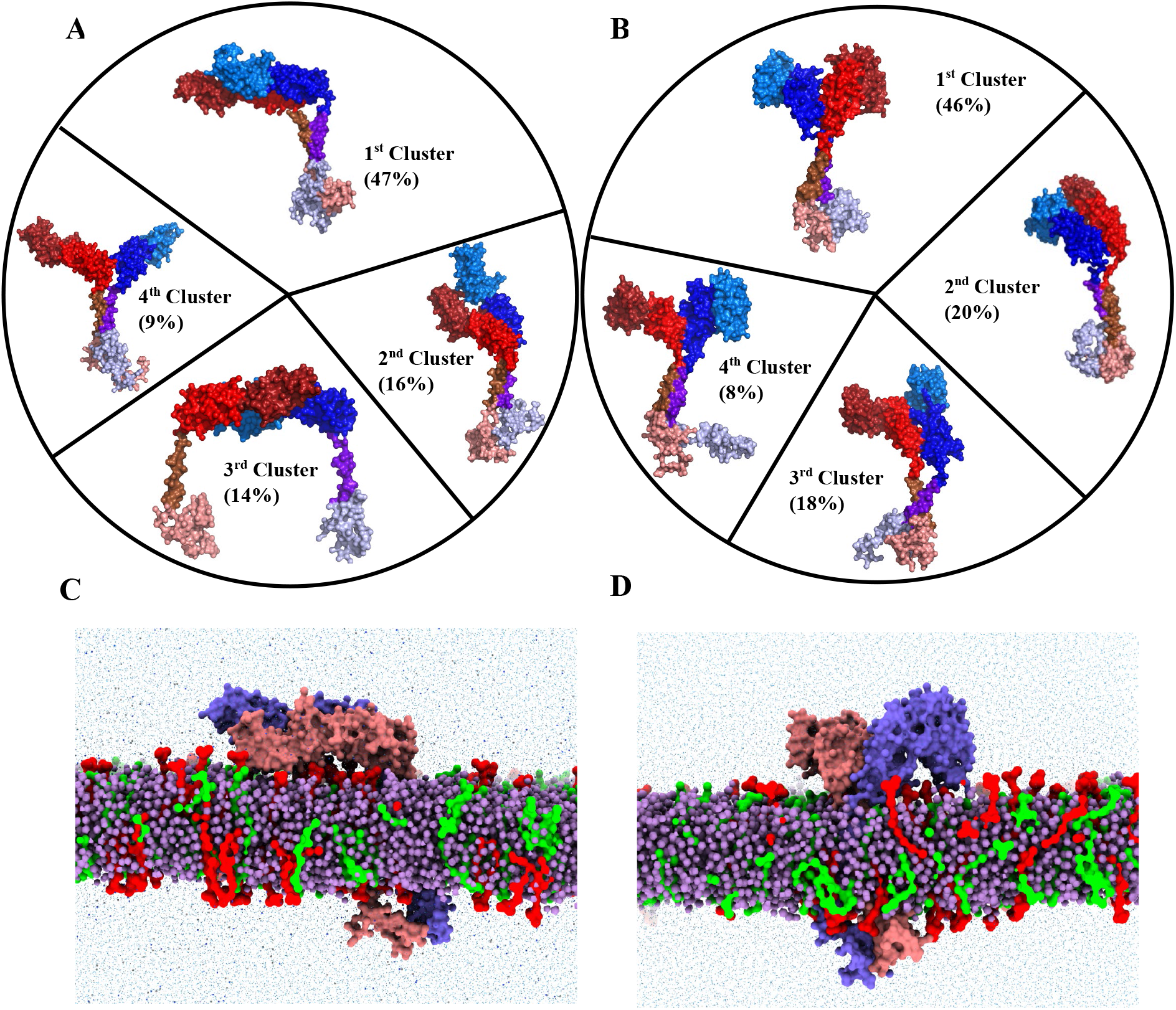
Comparison of the most populated clusters of the FN1-FN2-TM-JM construct dimers of EphA1 (A) and EphA2 (B). All conformations are shown in surface representation. The two monomers are designated as chain A and chain B throughout. In chain A, the FN1, FN2, TM, and JM regions are colored raspberry, red, chocolate, and salmon, respectively; in chain B, they are colored marine blue, blue, purple-blue, and light blue. Clustering of both EphA1 and EphA2 dimers was performed using a 2nm cutoff based on the final 2.5 µs of each trajectory from eight simulations. (C, D) Representative snapshots of the dominant population clusters of the FN1-FN2-TM-JM construct dimers of EphA1 (C) and EphA2 (D). Dimers are shown in surface representation, with chain A and chain B displayed as orange-red and purple surfaces, respectively. PIP2 and POPS are shown as red and green spheres, respectively, while POPC is shown as purple spheres.

Comparative mapping of FN-domain interfacial residues (Fig. S11) further distinguished the receptors. EphA1 displayed very distributed intermolecular contacts across both FN1 and FN2 domains, producing relatively symmetric docking geometries. This is also true when we examine to contact map of the FN2 domain in the shorter single FN domain dimer (Fig. S12). EphA2, in contrast, showed an asymmetric configuration in which the FN2 domain of chain B interacted simultaneously with both FN1 and FN2 of chain A, an arrangement facilitated by diverse conformational states of the linker between the domains. Similarly, even the interaction of the single FN domains in the FN2-TM-JM protein is asymmetric, with the N-termini of the FN2 domain mostly pointing away from one another. This is amplified in the tandem FN protein, placing FN1 domains away from one another, especially in the EphA2 case where the linker is more kinked. Together, these results indicate that extracellular FN domain positioning and their particle interactions, JM-PIP2 engagement, and lipid interactions collectively modulate TM dimer geometry and stability, providing a mechanism by which Eph receptors fine-tune signaling in response to membrane composition and extracellular architecture.

### FN1-FN2 linker as a structural basis for differential signaling

To better understand the configurational flexibility between the FN1 and FN2 domains particularly in EphA2, we compared FN1-FN2 arrangements from available crystal structures of the extracellular domains of ligand-bound and unbound EphA2, EphA4, and EphB2 (Fig. 4). In EphA2, the FN1 domain in ligand-bound configurations is positioned closer to FN2 compared to the unbound form (Fig. 4A), indicating that ligand engagement can induce or stabilize interdomain closure. This observation is consistent with previous reports showing that Eph receptor activation involves ligand-induced rearrangements of the ectodomain (including FN domains), which propagate configurational changes via the TM toward the membrane-proximal juxtamembrane regions, ultimately modulating receptor clustering and downstream signaling.^43–45^ The FN1-FN2 model of EphA1 used here, derived from unbound EphA2 crystal structure, adopts a similarly open configuration (Fig. 4B). Structures of EphA4 and EphB2 show even more pronounced FN1-FN2 closure upon ligand binding, underscoring the potential of the interdomain linker to function as a hinge that modulates relative domain orientations (Fig. 4C). To explore the basis for these differences, we compared the linker sequences of EphA1, EphA2, EphA4, and EphB2 (Fig. 4D). The linkers of EphA2, EphA4, and EphB2 are enriched in polar uncharged residues (Ser, Asn, Gln, Thr) and terminate with a proline. This composition is consistent with higher intrinsic disorder, as polar residues promote flexibility while proline can introduce kinks and disrupt regular secondary structure.^46^ In contrast, the EphA1 linker contains a methionine (M) and histidine (H) residue with bulkier, less conformationally dynamic side chains which could impart rigidity and restrict interdomain motion. We also examined the conservation of the FN1-FN2 region to align the amino acid sequences of EphA1 and A2 receptors across species (Fig. S13). Interestingly, only Ile 440 and Met 442 are highly conserved in the EphA1 linker region whereas in EphA2 the level of conservation is much higher. This includes two Prolines at the C-terminus of the linker which likely stabilize different orientation states. These differences are reflected in our simulations, where EphA2 consistently exhibited greater FN1-FN2 flexibility than EphA1.

**Figure 4.**
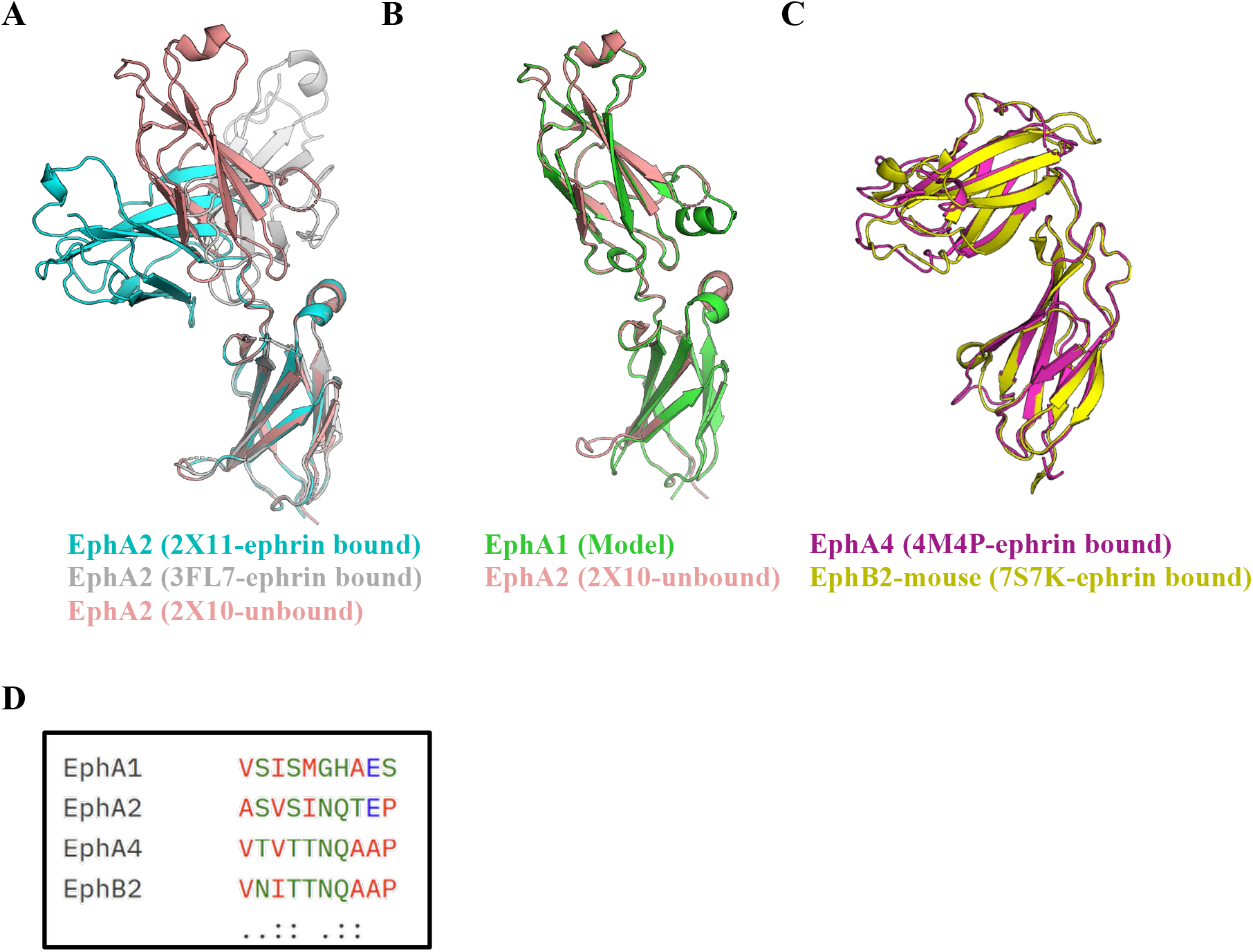
Structural alignment of the FN1-FN2 regions from the crystal structures: (A) Ligand bound and unbound conformations of EphA2, (B) The homology model of EphA1 based on the unbound structure of EphA2 and (C) Bound conformations for both EphA4 and EphB2. (D) Sequence alignment of the linker region between the FN1 and FN2 domains of all the four receptors.

We quantified interdomain flexibility by measuring the rotation angle between FN1 and FN2 throughout the simulations (Fig. 5). EphA1 maintained a narrow angle distribution (∼150° to 160°), indicative of a rigid, extended conformation. EphA2 displayed a broader distribution (∼90° to 150°), reflecting substantial bending and conformational variability. This structural plasticity may underlie EphA2’s capacity for ligand-independent signaling: a flexible FN1-FN2 hinge allows the extracellular domains to adopt multiple configurations, facilitating transient receptor-receptor contacts, pre-formed dimers or oligomers, and interactions with membrane lipids or co-receptors.^43,45,47^ EphA2 also engages in homotypic head-to-tail interactions between FN and LBD domains (L-shaped dimers) in the absence of ligand, further contributing to basal signaling activity. By contrast, the rigid FN1-FN2 architecture of EphA1 necessitates ligand engagement to achieve the proper TM orientation for stable dimerization and downstream signaling. The combination of a flexible FN1-FN2 linker in EphA2 and a rigid linker in EphA1 thus establishes a mechanistic basis for their functional divergence, with EphA2’s configurational adaptability enabling broader signaling modalities, ligand-independent activity, and sensitivity to membrane composition, mechanical forces, or heterotypic interactions.

**Figure 5.**
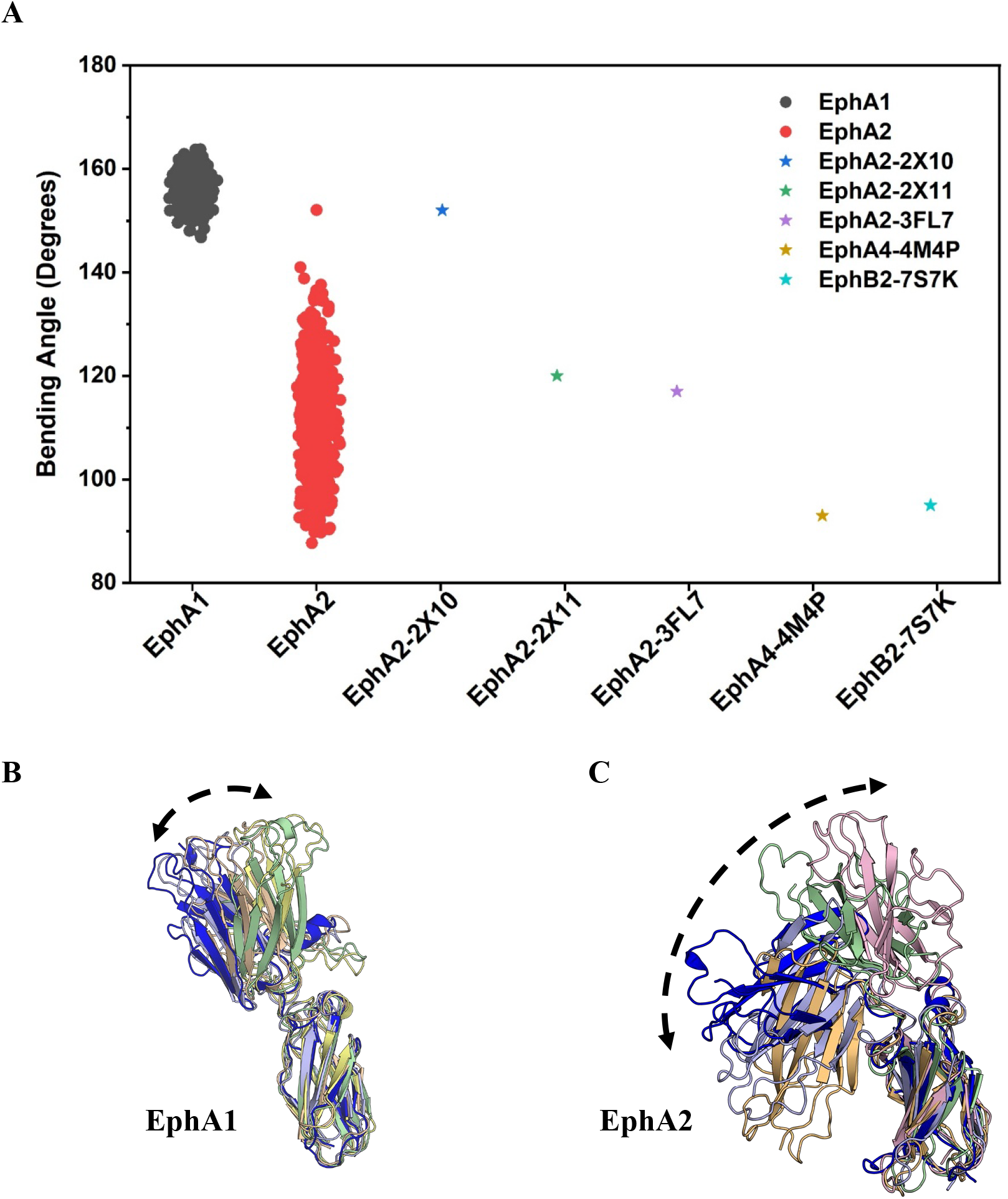
Comparison of the bending angle between the FN1 and FN2 domains evolved from the simulations for EphA1 (A) and EphA2 (B) with the available crystal structures. Cartoon representations of the FN1-FN2 bending from the ensemble of conformations observed from CG simulations. This correspondence is remarkable because the crystal structures do not include FN-FN domain contacts/dimers, suggesting that the differences are due to primary structure (Fig. 4).

## Discussion

The goal of comparing constructs of different lengths, consisting of regions which correspond to the fragments generated by proteases, was to examine their behaviour, indicating their likely physiological function but also to dissect how these membrane-adjacent extracellular domains and the juxtamembrane (JM) segments influence the dimerization behavior of EphA1 and EphA2. Although the solution NMR structures of the isolated TM domains have defined helix-helix interfaces apparently relevant to activation, these studies do not suggest how the geometry, linker flexibility and lipid interactions of these FN and JM domains modulate dimer formation at the membrane.^39,40^ Prior studies emphasize the importance of the ligand binding and cysteine rich domains (LBD and CRD, respectively) in ligand-independent dimerization;^18^ however, these domains are far from the membrane and it has remained unclear whether the more membrane-proximal FN1 and FN2 domains as well as the JM region can also exert regulatory effects. Our approach comparing systems of different protein lengths and domain complexity allowed us to explore how these structural layers integrate to tune dimerization in receptor specific ways.

A central outcome of this study is that that while both EphA1 and EphA2 TM segments possess an inherent ability to dimerize in anionic lipid membranes (shown by us and others), the modes of association differ markedly between the two receptors. EphA1 preferentially adopts symmetric AXXXGXXXG-centered dimers, whereas EphA2 samples a broader ensemble, including leucine zipper–like interfaces. The emergence of such polymorphism in EphA2 suggests a higher degree of configurational variability, a feature also observed in RTKs such as EGFR and ErbB2 where multiple TM states are proposed to underlie transitions between inactive, intermediate, and active structures.^28,48–51^ By analogy, the heterogeneity in EphA2 TM packing may facilitate a dynamic equilibrium between signaling states, supporting its well-documented ability to participate in ligand-independent signaling and receptor cross-talk. EphA1, conversely, appears more structurally constrained, consistent with a receptor that relies more heavily on canonical, ligand-driven dimerization.

The behavior of the FN domains provides additional mechanistic insight. The strong inhibition of TM association in the FN2-TM-JM constructs indicates that membrane proximal extracellular mass and orientation can directly restrict TM helix proximity. An upright FN2 domain acts as a steric and orientational barrier, suggesting that the Eph receptors likely use their extracellular architecture as a regulatory gate controlling when TM association is permitted and when it is sterically disfavored. Strikingly, the addition of the second FN (FN1) domain reverses this inhibition, restoring stable dimerization in both EphA1 and EphA2. This indicates that FN1 reshapes the orientation of FN2, either by altering its tilt relative to the membrane or by stabilizing configurations in which FN2 no longer sterically blocks TM engagement. This interplay between FN1 and FN2 echoes the more general principle that multi-domain extracellular architectures function not only as ligand-binding platforms but also as mechanical and geometric scaffolds that predispose receptors toward particular transmembrane configurations.^52,53^ Residue-residue contact maps between the FN regions (Fig. S11) further revealed that these extracellular docking arrangements differ substantially between the two receptors. EphA1 forms broadly distributed intermolecular contacts involving both the FN1 and FN2 domains, resulting in a relatively symmetric interface. In contrast, EphA2 adopts a more asymmetric docking mode in which the FN2 domain of one protomer interacts simultaneously with both the FN1 and FN2 domains of the opposing protomer. These distinct interface organizations are consistent with the greater conformational flexibility of the EphA2 FN1-FN2 linker, which enables a wider range of extracellular domain orientations.

Importantly in interactions with the membrane surface, EphA1 exhibits more extensive FN-PIP₂ contacts and stronger PIP₂ clustering, whereas EphA2 shows more selective and asymmetric interactions. These differences may help tune the local lipid environment during receptor assembly. Interestingly, the influence of lipid composition on EphA2 aligns with our earlier observation that cholesterol promotes both left-handed and right-handed TM dimer formations,^51^ highlighting that EphA2 remains highly responsive to changes in membrane biophysical properties as well as to mechanical cues or interactions with other membrane proteins.^54,55^ Moreover, because PIP₂ is known to regulate clustering, cooperativity, and activation thresholds in several RTKs, the stronger FN-PIP₂ engagement in EphA1 may contribute to more stable, ligand-dependent assemblies, while the weaker and asymmetric engagement in EphA2 may support dynamic, lipid-sensitive signaling states. The receptor-specific differences in FN1/FN2 interactions with the membrane, particularly with anionic lipids, further suggest that the extracellular domains can respond to changes in the membrane environment. It should be noted, however, that the membrane model employed in this study does not reproduce the physiological lipid asymmetry of healthy mammalian plasma membranes, where PIP₂ is predominantly localized to the cytoplasmic leaflet while the outer leaflet is largely composed of zwitterionic lipids.^56^ Therefore, the observed interactions between the FN domains and PIP₂ should be interpreted as mechanistic observations within the chosen membrane model rather than direct representations of healthy cell membranes. Nevertheless, disruption of plasma membrane lipid asymmetry is a well-established feature of many pathological conditions, including cancer, where increased exposure of anionic lipids on the extracellular leaflet has been reported. Under such conditions, extracellular lipid interactions may contribute more significantly to receptor organization and membrane association. Accordingly, our objective was not to reproduce the membrane composition of a specific cell type, but to investigate how an anionic membrane environment influences receptor organization and lipid interactions under well-controlled conditions. Similar simplified membrane models have been widely used in Martini simulations of receptor tyrosine kinases to dissect lipid-dependent effects.^51,57,58^ While such models do not capture the full complexity of native plasma membranes, they provide a useful framework for identifying receptor-specific membrane interactions and conformational behavior.

The flexibility of the FN1-FN2 linker emerges as another key determinant underlying many of these behaviors, as already envisaged in earlier work.^34^ EphA2’s polar, proline-terminated linker allows hinge-like bending and greater reorientation freedom of FN2 relative to FN1, enabling the extracellular region to adapt to membrane curvature, lipid microdomains, or neighboring proteins. This adaptability provides a mechanistic explanation for EphA2’s ability to engage diverse partners and participate in ligand-independent signaling. In contrast, the more rigid EphA1 linker constrains the motion of FN2, enforcing a more fixed extracellular arrangement that may bias the receptor toward specific TM associations and a more classical ligand-triggered activation mode. This structural rigidity aligns with experimental observations that EphA1 signaling is more dependent on ligand engagement and less prone to ligand-independent activation than EphA2. In particular, an L-shaped arrangement has been inferred for ligand independent EphA2 and is seen in the x-ray structure of EphA4 where the FN domains serve as a second ligand binding domain interacting with Ephrin in cis or the LBD of another receptor unit that lies down adjacently on the membrane.^59–62^ Intriguingly, an alignment of EphA1 and A2 sequences over many species (Fig. S13) not only points to differences in the linker region, mentioned above, but also shows that overall EphA2 sequences are more conserved especially in the membrane proximal FN2 region than for EphA1. At first glance these findings may appear contradictory: should not fewer specific interactions lead to more flexibility? We argue (also with reference to Fig. S11) that the opposite is the case: because the FN-FN domain interactions are weaker and more spread out in EphA1, the linker experiences less strain and varies less in orientation. While this argument reflects on strain and frustration in protein structures^63^ the greater sequence conservation in EphA2 FN regions and in the linker would be consistent with the evolution of specific interactions, between the FN domains, with other proteins and more distinct lipids for EphA2.

Lipid interactions at the intracellular side reinforce these themes. The JM region is heavily enriched in basic residues known to interact with anionic lipids such as PIP₂. Our comparison between constructs demonstrates that JM-PIP₂ interactions are substantially enhanced in the presence of both FN domains, for both receptors. These interactions are coordinated, multi-layered mechanism where extracellular orientation influences how the intracellular JM approaches the membrane, which in turn modulates lipid recruitment and TM association. The finding that FN containing constructs exhibit increased JM-PIP₂ contacts suggests that extracellular and intracellular domains operate in concert to stabilize or destabilize TM packing, supporting a model in which Eph receptors integrate signals across their entire length, and possibly by preferred lipid-type clustering in the bilayer^51^ rather than functioning as modular, independently behaving segments.

We noted limitations of AlphFold3 in the introduction in not being able accommodate features of the lipid bilayer and yield good reliability scores for predictions in some systems. We found in this study that the scores for EphA1 FN-TM-JM and FN-FN-M-JM dimers are particularly low (Fig. S14A-B), emphasizing the role of the membrane in stabilizing certain configurations of the domains. For EphA2 these scores are slightly higher at a-b (Fig. S14C-D). It should be noted that EphA2 crystal structures of the entire extracellular receptor region (also determined in absence of lipids or lipid mimicking detergent) do not contain FN-FN domain contacts. Another limitation of this study is that the receptor constructs were simulated without extracellular glycosylation. Glycosylation of the FN1 domain and the FN1-FN2 linker may influence linker flexibility and extracellular domain organization. Future simulations incorporating experimentally characterized glycosylation patterns will help assess these effects.

Several limitations of the coarse-grained simulations must be acknowledged. As discussed in our 2023 study,^28^ CG force fields may under-stabilize certain packing motifs and cannot resolve side-chain level rearrangements that provide sufficiently high steric barriers to prevent switching between TM interfaces, often leading to TM dimer asymmetry due to helix sliding and/or rotation. Nonetheless, the trends observed correspond well with primary-sequence differences in the TM helix, JM region, and extracellular linkers, lending confidence to the mechanistic framework. Moreover, the use of a POPC/POPS/PIP₂ membrane provided a well-defined anionic membrane environment, rather than the DMPC bilayers used in earlier studies, provide lipid-driven effects that align more closely with known biophysical behavior of Eph receptors. These insights set the stage for testable hypotheses involving mutagenesis of linker residues, targeted alteration of FN orientations, and lipid composition experiments in reconstituted or cellular systems.

Taken together, these findings support a model in which EphA1 and EphA2 differ not simply in TM interface preference but as an integrated, domain-spanning regulatory system. FN orientation, linker flexibility, and lipid engagement collectively shape the configurational landscape accessible to each receptor. EphA1 behaves as a structurally constrained receptor optimized for stable, ligand-dependent assembly, whereas EphA2 maintains adaptable extracellular and TM configurations capable of responding to diverse extracellular cues and membrane environments. These mechanisms provide a structural rationale for their divergent physiological and pathological roles and underscore the importance of considering the full receptor architecture not only the TM segment when developing therapeutic strategies targeting Eph receptors.

## Methods

### Modeling of the EphA1 & EphA2 constructs

The sequences for human EphA1 and EphA2 modeling were extracted from the UniProt database (UniProtKB: P21709 and P29317). We modeled three different fragments of both EphA1 and EphA2 (shown schematically with sequence limits in Fig S1B). The smallest constructs include the transmembrane (TM) region with short N- and C-terminal extensions. The TM domains were modeled as an ideal α-helix using Discovery studio Visualizer v24 and we added an extra 8-9 residues at the N-terminal and 10 residues at the C-terminal side of the TM in an extended conformation. The next level constructs include the TM region with the 2^nd^ domain of fibronectin III (FN2) on the N-terminus and the complete juxtamembrane (JM) region at their C-terminus. Our largest constructs include both the 1^st^ and the 2^nd^ FN III (FN1 and FN2) domains on the N-terminus and the JM region on the C-terminus. The structure of the FN1 and FN2 domains of EphA2 were extracted from the available ectodomain X-ray crystal structure (PDB ID: 2X10) and the missing regions were modeled using Modeller.^64^ For EphA1, the FN1 and FN2 domains were modeled based on the ectodomain structure of EphA2 using the homology modeling approach in the SWISS-MODEL server.^65^ The JM regions of both EphA1 and EphA2 were modeled in the initial structures in the extended conformation because JM regions are highly flexible and unstructured.

### Coarse-grained molecular dynamics simulation

To check the homodimerization for each construct, we built a system with the construct monomers placed 5.0 nm apart from each other (Fig S1C). The atomistic (AT) modeled systems of all the constructs of EphA1 and EphA2 were converted to coarse-grained (CG) representation using the *martinize2.py* workflow module of the MARTINI 3 force field considering the secondary structure DSSP assignment.^66^ CG simulations were performed using Gromacs version 2023.3.^67^ The *insane.py* script^68^ was used for the setting up of the lipid bilayer. The lipid bilayer is composed of POPC (80%), POPS (15%) and PIP2 (5%) evenly distributed in both the upper and lower leaflet. The membrane composition was selected to provide a simplified and reproducible model for investigating receptor-lipid interactions. Because plasma membrane lipid composition varies substantially between cell types and comprehensive lipidomic data are not available for EphA1- or EphA2-expressing cells, we did not attempt to model a specific cellular membrane. Instead, the same membrane composition was used for all systems to enable direct comparison of receptor behavior under identical lipid conditions. For the smaller constructs, the monomers are placed in a cubic box of dimensions 10×10×10 nm^3^ with around 248 POPC, 46 POPS, 14 PIP2 lipids and 4500 CG water molecules. For the medium-size constructs, the box dimensions were 15×15×15 nm^3^ with about 548 POPC, 102 POPS, 34 PIP2 and 18700 CG water molecules. For the largest constructs of EphA1 and EphA2, the box size was increased to 30×30×30 nm^3^ accommodating about 2395 POPC, 448 POPS, 148 PIP2 and 183000 CG water molecules. The pH of the systems was considered neutral. All the simulations were run in presence of regular Martini CG water and neutralised and ions added to 0.15M NaCl. The systems were equilibrated for 500 ps. The long-range electrostatic interactions were used with reaction field type having a cutoff value of 1.1 nm. We used potential-shift-verlet for the Lennard-Jones interactions with a value of 1.1 nm for the cutoff scheme. V-rescale thermostat with a reference temperature of 320 K in combination with a Berendsen barostat with a coupling constant of 1.0 ps, compressibility of 3.0 × 10^−4^ bar ^−1^, and a reference pressure of 1 bar was used. The integration time step was 20 fs. A temperature of 320 K was chosen to modestly enhance conformational sampling within the accessible simulation timescale while maintaining the structural integrity of the receptor systems. The simulations of the smaller constructs were run in quadruplicate for 4 µs each whereas for the medium and the larger constructs we ran 8 replica simulations for 4 µs each.

### Data Analysis

Analysis of the trajectories was carried out by implementing the modules built into Gromacs. The contact maps between the helices and the FN domains were calculated with a distance cut off 0.55 nm and 0.9 nm for all the backbone and side-chain atoms, respectively. Data were plotted in Origin2024b. AlphaFold3 webserver^63^ was also used to predict the homodimer structures. We used the WebLogo server^69^ to align EphA1 and A2 sequences across as many species as the program deemed reasonable. Different cutoffs were selected for clustering of the different constructs empirically after visual inspection of the clustering results to provide a meaningful separation of the dominant conformational states while avoiding excessive fragmentation of structurally similar conformations.

## Author Contributions

ARS and MB designed the project. ARS generated the protein coarse-grained models and performed the MD simulations and analysed the data. NB provided additional advice throughout. ARS and MB co-wrote the paper. All authors read and approved the final version of the manuscript and supporting information.

## Acknowledgements

This work utilized the high-performance computing resources at CWRU and the Buck lab. was supported by NIH grants R21AG084065, R01EY029169 when the project began and is currently supported by R01AG089561.

## Data and Software Availability Statement

All input files necessary to reproduce our CG simulations are freely available at https://github.com/amita-bucklab/EphA1_vs_EphA2-dimerization. Molecular dynamics simulations were performed using the GROMACS 2023 package, with detailed protocol reported in the Materials and Methods section of the manuscript. Plotting was performed using Origin2024b, and structural rendering was done with PyMOL and ChimeraX.

## Supporting Information

Supporting figures are included in the supporting information.

## References

(1) Ma, X.; Yu, H. Global Burden of Cancer. Yale J. Biol. Med. 2006, 79 (3–4), 85–94.

(2) Global, Regional, and National Life Expectancy, All-Cause Mortality, and Cause-Specific Mortality for 249 Causes of Death, 1980-2015: A Systematic Analysis for the Global Burden of Disease Study 2015. Lancet (London, England) 2016, 388 (10053), 1459–1544. 10.1016/S0140-6736(16)31012-1.

(3) Schnerch, D.; Yalcintepe, J.; Schmidts, A.; Becker, H.; Follo, M.; Engelhardt, M.; Wäsch, R. Cell Cycle Control in Acute Myeloid Leukemia. Am. J. Cancer Res. 2012, 2 (5), 508–528.

(4) Masson, N.; Ratcliffe, P. J. Hypoxia Signaling Pathways in Cancer Metabolism: The Importance of Co-Selecting Interconnected Physiological Pathways. Cancer Metab. 2014, 2 (1), 3. 10.1186/2049-3002-2-3.

(5) Abbaspour Babaei, M.; Kamalidehghan, B.; Saleem, M.; Huri, H. Z.; Ahmadipour, F. Receptor Tyrosine Kinase (c-Kit) Inhibitors: A Potential Therapeutic Target in Cancer Cells. Drug Des. Devel. Ther. 2016, 10, 2443–2459. 10.2147/DDDT.S89114.

(6) Regad, T. Targeting RTK Signaling Pathways in Cancer. Cancers (Basel). 2015, 7 (3), 1758–1784. 10.3390/cancers7030860.

(7) Buckens, O. J.; El Hassouni, B.; Giovannetti, E.; Peters, G. J. The Role of Eph Receptors in Cancer and How to Target Them: Novel Approaches in Cancer Treatment. Expert Opin. Investig. Drugs 2020, 29 (6), 567–582. 10.1080/13543784.2020.1762566.

(8) Pasquale, E. B. Eph Receptors and Ephrins in Cancer: Bidirectional Signalling and Beyond. Nat. Rev. Cancer 2010, 10 (3), 165–180. 10.1038/nrc2806.

(9) Barquilla, A.; Pasquale, E. B. Eph Receptors and Ephrins: Therapeutic Opportunities. Annu. Rev. Pharmacol. Toxicol. 2015, 55, 465–487. 10.1146/annurev-pharmtox-011112-140226.

(10) Hafner, C.; Becker, B.; Landthaler, M.; Vogt, T. Expression Profile of Eph Receptors and Ephrin Ligands in Human Skin and Downregulation of EphA1 in Nonmelanoma Skin Cancer. Mod. Pathol. an Off. J. United States Can. Acad. Pathol. Inc 2006, 19 (10), 1369–1377. 10.1038/modpathol.3800660.

(11) Herath, N. I.; Doecke, J.; Spanevello, M. D.; Leggett, B. A.; Boyd, A. W. Epigenetic Silencing of EphA1 Expression in Colorectal Cancer Is Correlated with Poor Survival. Br. J. Cancer 2009, 100 (7), 1095–1102. 10.1038/sj.bjc.6604970.

(12) Biao-xue, R.; Xi-guang, C.; Shuan-ying, Y.; Wei, L.; Zong-juan, M. EphA2-Dependent Molecular Targeting Therapy for Malignant Tumors. Curr. Cancer Drug Targets 2011, 11 (9), 1082–1097. 10.2174/156800911798073050.

(13) Wykosky, J.; Debinski, W. The EphA2 Receptor and EphrinA1 Ligand in Solid Tumors: Function and Therapeutic Targeting. Mol. Cancer Res. 2008, 6 (12), 1795–1806. 10.1158/1541-7786.MCR-08-0244.

(14) Xiao, T.; Xiao, Y.; Wang, W.; Tang, Y. Y.; Xiao, Z.; Su, M. Targeting EphA2 in Cancer. J. Hematol. Oncol. 2020, 13 (1), 114. 10.1186/s13045-020-00944-9.

(15) Ieguchi, K.; Maru, Y. Roles of EphA1/A2 and Ephrin-A1 in Cancer. Cancer Sci. 2019, 110 (3), 841–848. 10.1111/cas.13942.

(16) Janes, P. W.; Nievergall, E.; Lackmann, M. Concepts and Consequences of Eph Receptor Clustering. Semin. Cell Dev. Biol. 2012, 23 (1), 43–50. 10.1016/j.semcdb.2012.01.001.

(17) Pasquale, E. B. Eph-Ephrin Bidirectional Signaling in Physiology and Disease. Cell 2008, 133 (1), 38–52. 10.1016/j.cell.2008.03.011.

(18) Liang, L.-Y.; Patel, O.; Janes, P. W.; Murphy, J. M.; Lucet, I. S. Eph Receptor Signalling: From Catalytic to Non-Catalytic Functions. Oncogene 2019, 38 (39), 6567–6584. 10.1038/s41388-019-0931-2.

(19) Matsumoto, M.; Gomez-Soler, M.; Lombardi, S.; Lechtenberg, B. C.; Pasquale, E. B. Missense Mutations of the Ephrin Receptor EPHA1 Associated with Alzheimer&#x2019;s Disease Disrupt Receptor Signaling Functions. J. Biol. Chem. 2025, 301 (2). 10.1016/j.jbc.2024.108099.

(20) Kim, Y.; Lasso, G.; Patel, H.; Vardarajan, B.; Santa-Maria, I.; Lefort, R. Alzheimer’s Disease-Associated P460L Mutation in Ephrin Receptor Type A1 (EphA1) Leads to Dysregulated Rho-GTPase Signaling. bioRxiv 2021, 2021.06.17.448790. 10.1101/2021.06.17.448790.

(21) Owens, H. A.; Thorburn, L. E.; Walsby, E.; Moon, O. R.; Rizkallah, P.; Sherwani, S.; Tinsley, C. L.; Rogers, L.; Cerutti, C.; Ridley, A. J.; Williams, J.; Knäuper, V.; Ager, A. Alzheimer’s Disease-Associated P460L Variant of EphA1 Dysregulates Receptor Activity and Blood-Brain Barrier Function. Alzheimers. Dement. 2024, 20 (3), 2016–2033. 10.1002/alz.13603.

(22) Gatto, G.; Morales, D.; Kania, A.; Klein, R. EphA4 Receptor Shedding Regulates Spinal Motor Axon Guidance. Curr. Biol. 2014, 24 (20), 2355–2365. 10.1016/j.cub.2014.08.028.

(23) Eriksson, O.; Thulin, Å.; Asplund, A.; Hegde, G.; Navani, S.; Siegbahn, A. Cross-Talk between the Tissue Factor/Coagulation Factor VIIa Complex and the Tyrosine Kinase Receptor EphA2 in Cancer. BMC Cancer 2016, 16 (1), 341. 10.1186/s12885-016-2375-1.

(24) Kikuchi, K.; Kozuka-Hata, H.; Oyama, M.; Seiki, M.; Koshikawa, N. Identification of Proteolytic Cleavage Sites of EphA2 by Membrane Type 1 Matrix Metalloproteinase on the Surface of Cancer Cells. Methods Mol. Biol. 2018, 1731, 29–37. 10.1007/978-1-4939-7595-2_3.

(25) Eriksson, O.; Ramström, M.; Hörnaeus, K.; Bergquist, J.; Mokhtari, D.; Siegbahn, A. The Eph Tyrosine Kinase Receptors EphB2 and EphA2 Are Novel Proteolytic Substrates of Tissue Factor/Coagulation Factor VIIa. J. Biol. Chem. 2014, 289 (47), 32379–32391. 10.1074/jbc.M114.599332.

(26) Bessman, N. J.; Freed, D. M.; Lemmon, M. A. Putting Together Structures of Epidermal Growth Factor Receptors. Curr. Opin. Struct. Biol. 2014, 29, 95–101. 10.1016/j.sbi.2014.10.002.

(27) Abramson, J.; Adler, J.; Dunger, J.; Evans, R.; Green, T.; Pritzel, A.; Ronneberger, O.; Willmore, L.; Ballard, A. J.; Bambrick, J.; Bodenstein, S. W.; Evans, D. A.; Hung, C.-C.; O’Neill, M.; Reiman, D.; Tunyasuvunakool, K.; Wu, Z.; Žemgulytė, A.; Arvaniti, E.; Beattie, C.; Bertolli, O.; Bridgland, A.; Cherepanov, A.; Congreve, M.; Cowen-Rivers, A. I.; Cowie, A.; Figurnov, M.; Fuchs, F. B.; Gladman, H.; Jain, R.; Khan, Y. A.; Low, C. M. R.; Perlin, K.; Potapenko, A.; Savy, P.; Singh, S.; Stecula, A.; Thillaisundaram, A.; Tong, C.; Yakneen, S.; Zhong, E. D.; Zielinski, M.; Žídek, A.; Bapst, V.; Kohli, P.; Jaderberg, M.; Hassabis, D.; Jumper, J. M. Accurate Structure Prediction of Biomolecular Interactions with AlphaFold 3. Nature 2024, 630 (8016), 493–500. 10.1038/s41586-024-07487-w.

(28) Sahoo, A. R.; Souza, P. C. T.; Meng, Z.; Buck, M. Transmembrane Dimers of Type 1 Receptors Sample Alternate Configurations: MD Simulations Using Coarse Grain Martini 3 versus AlphaFold2 Multimer. Structure 2023, 31 (6), 735–745.e2. 10.1016/j.str.2023.03.014.

(29) Shrestha, P.; Sahoo, A. R.; Iannucci, M.; Willard, B.; Buck, M. Tyrosine Phosphorylation and the Inhibitory C-Terminal SAM Domain Moderately Affect Transient Interactions in a EphA2 Cytoplasmic Fragment in Solution: A Combined Experimental and Molecular Modeling Study. bioRxiv 2025, 2025.09.29.679228. 10.1101/2025.09.29.679228.

(30) Arkhipov, A.; Shan, Y.; Das, R.; Endres, N. F.; Eastwood, M. P.; Wemmer, D. E.; Kuriyan, J.; Shaw, D. E. Architecture and Membrane Interactions of the EGF Receptor. Cell 2013, 152 (3), 557–569. 10.1016/j.cell.2012.12.030.

(31) Kästner, J.; Loeffler, H. H.; Roberts, S. K.; Martin-Fernandez, M. L.; Winn, M. D. Ectodomain Orientation, Conformational Plasticity and Oligomerization of ErbB1 Receptors Investigated by Molecular Dynamics. J. Struct. Biol. 2009, 167 (2), 117–128. 10.1016/j.jsb.2009.04.007.

(32) Lelimousin, M.; Limongelli, V.; Sansom, M. S. P. Conformational Changes in the Epidermal Growth Factor Receptor: Role of the Transmembrane Domain Investigated by Coarse-Grained MetaDynamics Free Energy Calculations. J. Am. Chem. Soc. 2016, 138 (33), 10611–10622. 10.1021/jacs.6b05602.

(33) Chavent, M.; Chetwynd, A. P.; Stansfeld, P. J.; Sansom, M. S. P. Dimerization of the EphA1 Receptor Tyrosine Kinase Transmembrane Domain: Insights into the Mechanism of Receptor Activation. Biochemistry 2014, 53 (42), 6641–6652. 10.1021/bi500800x.

(34) Chavent, M.; Seiradake, E.; Jones, E. Y.; Sansom, M. S. P. Structures of the EphA2 Receptor at the Membrane: Role of Lipid Interactions. Structure 2016, 24 (2), 337–347. 10.1016/j.str.2015.11.008.

(35) Kaszuba, K.; Grzybek, M.; Orłowski, A.; Danne, R.; Róg, T.; Simons, K.; Coskun, Ü.; Vattulainen, I. N-Glycosylation as Determinant of Epidermal Growth Factor Receptor Conformation in Membranes. Proc. Natl. Acad. Sci. 2015, 112 (14), 4334–4339. 10.1073/pnas.1503262112.

(36) Roberts, S. K.; Tynan, C. J.; Winn, M.; Martin-Fernandez, M. L. Investigating Extracellular in Situ EGFR Structure and Conformational Changes Using FRET Microscopy. Biochem. Soc. Trans. 2012, 40 (1), 189–194. 10.1042/BST20110632.

(37) Martin-Perez, M.; Urdiroz-Urricelqui, U.; Bigas, C.; Benitah, S. A. The Role of Lipids in Cancer Progression and Metastasis. Cell Metab. 2022, 34 (11), 1675–1699. 10.1016/j.cmet.2022.09.023.

(38) Skotland, T.; Kavaliauskiene, S.; Sandvig, K. The Role of Lipid Species in Membranes and Cancer-Related Changes. Cancer Metastasis Rev. 2020, 39 (2), 343–360. 10.1007/s10555-020-09872-z.

(39) Bocharov, E. V; Mayzel, M. L.; Volynsky, P. E.; Goncharuk, M. V; Ermolyuk, Y. S.; Schulga, A. A.; Artemenko, E. O.; Efremov, R. G.; Arseniev, A. S. Spatial Structure and PH-Dependent Conformational Diversity of Dimeric Transmembrane Domain of the Receptor Tyrosine Kinase EphA1. J. Biol. Chem. 2008, 283 (43), 29385–29395. 10.1074/jbc.M803089200.

(40) Bocharov, E. V; Mayzel, M. L.; Volynsky, P. E.; Mineev, K. S.; Tkach, E. N.; Ermolyuk, Y. S.; Schulga, A. A.; Efremov, R. G.; Arseniev, A. S. Left-Handed Dimer of EphA2 Transmembrane Domain: Helix Packing Diversity among Receptor Tyrosine Kinases. Biophys. J. 2010, 98 (5), 881–889. 10.1016/j.bpj.2009.11.008.

(41) Sharonov, G. V; Bocharov, E. V; Kolosov, P. M.; Astapova, M. V; Arseniev, A. S.; Feofanov, A. V. Point Mutations in Dimerization Motifs of the Transmembrane Domain Stabilize Active or Inactive State of the EphA2 Receptor Tyrosine Kinase. J. Biol. Chem. 2014, 289 (21), 14955–14964. 10.1074/jbc.M114.558783.

(42) Wirth, D.; Özdemir, E.; Wimley, W. C.; Pasquale, E. B.; Hristova, K. Transmembrane Helix Interactions Regulate Oligomerization of the Receptor Tyrosine Kinase EphA2. J. Biol. Chem. 2024, 300 (7), 107441. 10.1016/j.jbc.2024.107441.

(43) Himanen, J. P.; Goldgur, Y.; Miao, H.; Myshkin, E.; Guo, H.; Buck, M.; Nguyen, M.; Rajashankar, K. R.; Wang, B.; Nikolov, D. B. Ligand Recognition by A-Class Eph Receptors: Crystal Structures of the EphA2 Ligand-Binding Domain and the EphA2/Ephrin-A1 Complex. EMBO Rep. 2009, 10 (7), 722–728. 10.1038/embor.2009.91.

(44) Himanen, J. P.; Yermekbayeva, L.; Janes, P. W.; Walker, J. R.; Xu, K.; Atapattu, L.; Rajashankar, K. R.; Mensinga, A.; Lackmann, M.; Nikolov, D. B.; Dhe-Paganon, S. Architecture of Eph Receptor Clusters. Proc. Natl. Acad. Sci. U. S. A. 2010, 107 (24), 10860–10865. 10.1073/pnas.1004148107.

(45) Seiradake, E.; Harlos, K.; Sutton, G.; Aricescu, A. R.; Jones, E. Y. An Extracellular Steric Seeding Mechanism for Eph-Ephrin Signaling Platform Assembly. Nat. Struct. Mol. Biol. 2010, 17 (4), 398–402. 10.1038/nsmb.1782.

(46) Dyson, H. J.; Wright, P. E. Intrinsically Unstructured Proteins and Their Functions. Nat. Rev. Mol. Cell Biol. 2005, 6 (3), 197–208. 10.1038/nrm1589.

(47) Pasquale, E. B. Eph Receptor Signaling Complexes in the Plasma Membrane. Trends Biochem. Sci. 2024, 49 (12), 1079–1096. 10.1016/j.tibs.2024.10.002.

(48) Bocharov, E. V; Mineev, K. S.; Volynsky, P. E.; Ermolyuk, Y. S.; Tkach, E. N.; Sobol, A. G.; Chupin, V. V; Kirpichnikov, M. P.; Efremov, R. G.; Arseniev, A. S. Spatial Structure of the Dimeric Transmembrane Domain of the Growth Factor Receptor ErbB2 Presumably Corresponding to the Receptor Active State. J. Biol. Chem. 2008, 283 (11), 6950–6956. 10.1074/jbc.M709202200.

(49) Endres, N. F.; Das, R.; Smith, A. W.; Arkhipov, A.; Kovacs, E.; Huang, Y.; Pelton, J. G.; Shan, Y.; Shaw, D. E.; Wemmer, D. E.; Groves, J. T.; Kuriyan, J. Conformational Coupling across the Plasma Membrane in Activation of the EGF Receptor. Cell 2013, 152 (3), 543–556. 10.1016/j.cell.2012.12.032.

(50) Sahoo, A. R.; Buck, M. Structural and Functional Insights into the Transmembrane Domain Association of Eph Receptors. Int. J. Mol. Sci. 2021, 22 (16). 10.3390/ijms22168593.

(51) Sahoo, A. R.; Bhattarai, N.; Buck, M. Cholesterol-Dependent Dimerization and Conformational Dynamics of EphA2 Receptors from Coarse-Grained and All-Atom Simulations. Structure 2025, 33 (7), 1275–1287.e2. 10.1016/j.str.2025.03.014.

(52) Chataigner, L. M. P.; Leloup, N.; Janssen, B. J. C. Structural Perspectives on Extracellular Recognition and Conformational Changes of Several Type-I Transmembrane Receptors. Front. Mol. Biosci. 2020, Volume 7-2020.

(53) Valley, C. C.; Lewis, A. K.; Sachs, J. N. Piecing It Together: Unraveling the Elusive Structure-Function Relationship in Single-Pass Membrane Receptors. Biochim. Biophys. acta. Biomembr. 2017, 1859 (9 Pt A), 1398–1416. 10.1016/j.bbamem.2017.01.016.

(54) Singh, P. K.; Rybak, J. A.; Schuck, R. J.; Sahoo, A. R.; Buck, M.; Barrera, F. N.; Smith, A. W. Phosphatidylinositol 4,5-Bisphosphate Drives the Formation of EGFR and EphA2 Complexes. Sci. Adv. 2025, 10 (49), eadl0649. 10.1126/sciadv.adl0649.

(55) Schuck, R. J.; Ward, A. E.; Sahoo, A. R.; Rybak, J. A.; Pyron, R. J.; Trybala, T. N.; Simmons, T. B.; Baccile, J. A.; Sgouralis, I.; Buck, M.; Lamichhane, R.; Barrera, F. N. Cholesterol Inhibits Assembly and Oncogenic Activation of the EphA2 Receptor. Commun. Biol. 2025, 8 (1), 411. 10.1038/s42003-025-07786-6.

(56) Lorent, J. H.; Levental, K. R.; Ganesan, L.; Rivera-Longsworth, G.; Sezgin, E.; Doktorova, M.; Lyman, E.; Levental, I. Plasma Membranes Are Asymmetric in Lipid Unsaturation, Packing and Protein Shape. Nat. Chem. Biol. 2020, 16 (6), 644–652. 10.1038/s41589-020-0529-6.

(57) Hedger, G.; Sansom, M. S. P.; Koldsø, H. The Juxtamembrane Regions of Human Receptor Tyrosine Kinases Exhibit Conserved Interaction Sites with Anionic Lipids. Sci. Rep. 2015, 5 (1), 9198. 10.1038/srep09198.

(58) Abd Halim, K. B.; Koldsø, H.; Sansom, M. S. P. Interactions of the EGFR Juxtamembrane Domain with PIP2-Containing Lipid Bilayers: Insights from Multiscale Molecular Dynamics Simulations. Biochim. Biophys. Acta 2015, 1850 (5), 1017–1025. 10.1016/j.bbagen.2014.09.006.

(59) Kim, S.; Toth, P.; Wang, W.; Seiradake, E.; Shi, X.; Wang, B. EphA2 and Ephrin-A1 Utilize the Same Interface for Both in Cis and in Trans Interactions That Differentially Regulate Cell Signaling and Function. bioRxiv Prepr. Serv. Biol. 2025. 10.1101/2025.07.31.667925.

(60) Nikolov, D. B.; Xu, K.; Himanen, J. P. Eph/Ephrin Recognition and the Role of Eph/Ephrin Clusters in Signaling Initiation. Biochim. Biophys. Acta 2013, 1834 (10), 2160–2165. 10.1016/j.bbapap.2013.04.020.

(61) Shi, X.; Lingerak, R.; Herting, C. J.; Ge, Y.; Kim, S.; Toth, P.; Wang, W.; Brown, B. P.; Meiler, J.; Sossey-Alaoui, K.; Buck, M.; Himanen, J.; Hambardzumyan, D.; Nikolov, D. B.; Smith, A. W.; Wang, B. Time-Resolved Live-Cell Spectroscopy Reveals EphA2 Multimeric Assembly. Science 2023, 382 (6674), 1042–1050. 10.1126/science.adg5314.

(62) Xu, K.; Tzvetkova-Robev, D.; Xu, Y.; Goldgur, Y.; Chan, Y.-P.; Himanen, J. P.; Nikolov, D. B. Insights into Eph Receptor Tyrosine Kinase Activation from Crystal Structures of the EphA4 Ectodomain and Its Complex with Ephrin-A5. Proc. Natl. Acad. Sci. 2013, 110 (36), 14634 LP –14639. 10.1073/pnas.1311000110.

(63) Gianni, S.; Freiberger, M. I.; Jemth, P.; Ferreiro, D. U.; Wolynes, P. G.; Fuxreiter, M. Fuzziness and Frustration in the Energy Landscape of Protein Folding, Function, and Assembly. Acc. Chem. Res. 2021, 54 (5), 1251–1259. 10.1021/acs.accounts.0c00813.

(64) Fiser, A.; Sali, A. ModLoop: Automated Modeling of Loops in Protein Structures. Bioinformatics 2003, 19 (18), 2500–2501. 10.1093/bioinformatics/btg362.

(65) Waterhouse, A.; Bertoni, M.; Bienert, S.; Studer, G.; Tauriello, G.; Gumienny, R.; Heer, F. T.; de Beer, T. A. P.; Rempfer, C.; Bordoli, L.; Lepore, R.; Schwede, T. SWISS-MODEL: Homology Modelling of Protein Structures and Complexes. Nucleic Acids Res. 2018, 46 (W1), W296–W303. 10.1093/nar/gky427.

(66) Souza, P. C. T.; Alessandri, R.; Barnoud, J.; Thallmair, S.; Faustino, I.; Grünewald, F.; Patmanidis, I.; Abdizadeh, H.; Bruininks, B. M. H.; Wassenaar, T. A.; Kroon, P. C.; Melcr, J.; Nieto, V.; Corradi, V.; Khan, H. M.; Domański, J.; Javanainen, M.; Martinez-Seara, H.; Reuter, N.; Best, R. B.; Vattulainen, I.; Monticelli, L.; Periole, X.; Tieleman, D. P.; de Vries, A. H.; Marrink, S. J. Martini 3: A General Purpose Force Field for Coarse-Grained Molecular Dynamics. Nat. Methods 2021, 18 (4), 382–388. 10.1038/s41592-021-01098-3.

(67) Abraham, M. J.; Murtola, T.; Schulz, R.; Páll, S.; Smith, J. C.; Hess, B.; Lindahl, E. GROMACS: High Performance Molecular Simulations through Multi-Level Parallelism from Laptops to Supercomputers. SoftwareX 2015, 1-2, 19–25. 10.1016/j.softx.2015.06.001.

(68) Wassenaar, T. A.; Ingólfsson, H. I.; Böckmann, R. A.; Tieleman, D. P.; Marrink, S. J. Computational Lipidomics with Insane: A Versatile Tool for Generating Custom Membranes for Molecular Simulations. J. Chem. Theory Comput. 2015, 11 (5), 2144–2155. 10.1021/acs.jctc.5b00209.

(69) Crooks, G. E.; Hon, G.; Chandonia, J.-M.; Brenner, S. E. WebLogo: A Sequence Logo Generator. Genome Res. 2004, 14 (6), 1188–1190. 10.1101/gr.849004.

