## supplementary file for "Molecular Determinants of Receptor-Specific Membrane Interactions in EphA1 and EphA2 from Coarse-Grained Simulations"

**Figure S1.**

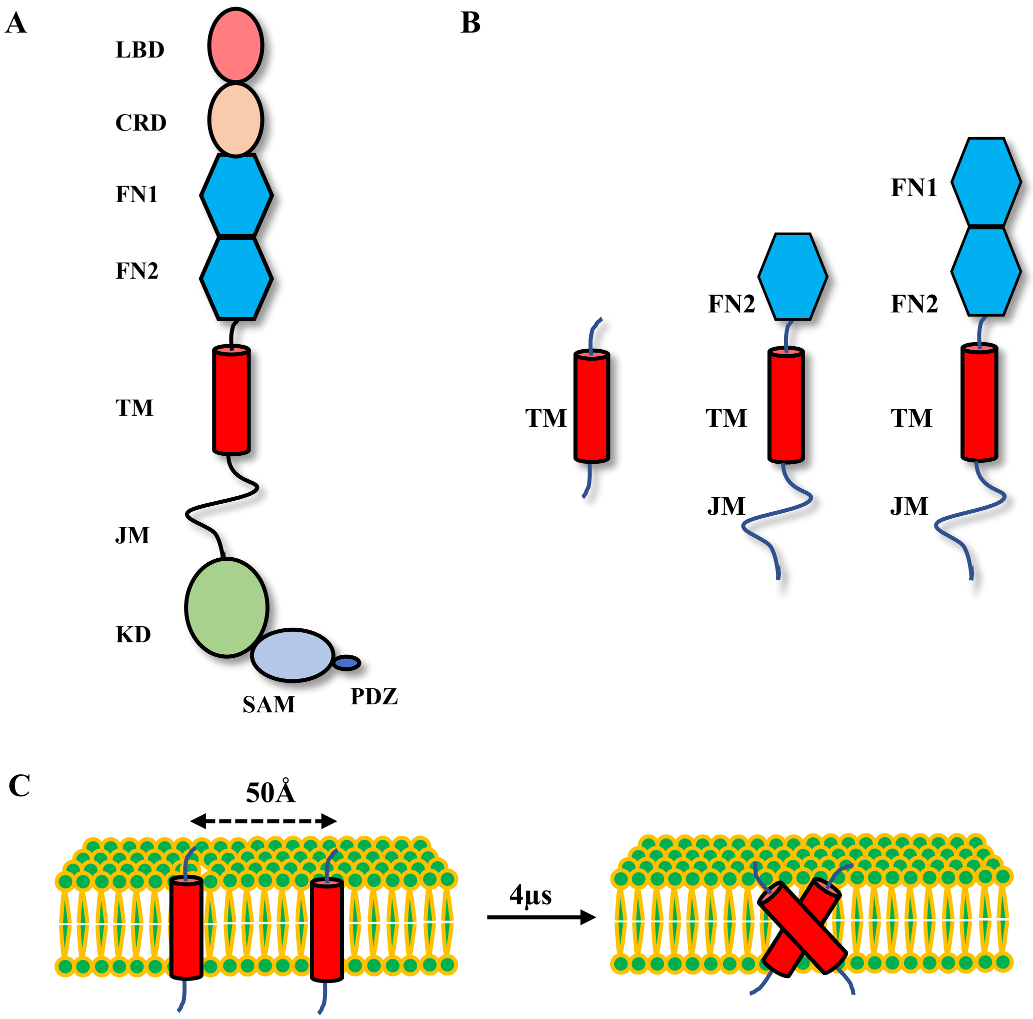


**Fig S1.** (A) Domain architecture of Eph receptors, showing the extracellular ligand-binding, cysteine-rich, and FNIII domains, a single-pass transmembrane helix, and the intracellular juxtamembrane, kinase, SAM, and PDZ-binding regions.. (B) Schematic representation of three different constructs of EphA1 and EphA2 used in this study. TM: Transmembrane region; FN2: 2$^{nd}$ Fibronectin domain; FN1: 1$^{st}$ Fibronectin domain; JM: Juxtamembrane region. Residues length for constructs: TM only (S540-Q578 for EphA1; E530-R568 for EphA2), FN2-TM-JM (G450-W623 for EphA1; N435-C612 for EphA2), FN1-FN2-TM-JM (G331-W623 for EphA1; R327-C612 for EphA2). (C) Association of dimers considering three constructs of EphA1 and EphA2 in the mixed lipid bilayer. Monomers for each system are initially placed 50Å apart from each other and then inserted into lipid bilayer and the simulations were extended upto 4μs to verify the dimerization and domains involved in the dimeric interface of these constructs.

#### Figure S2.

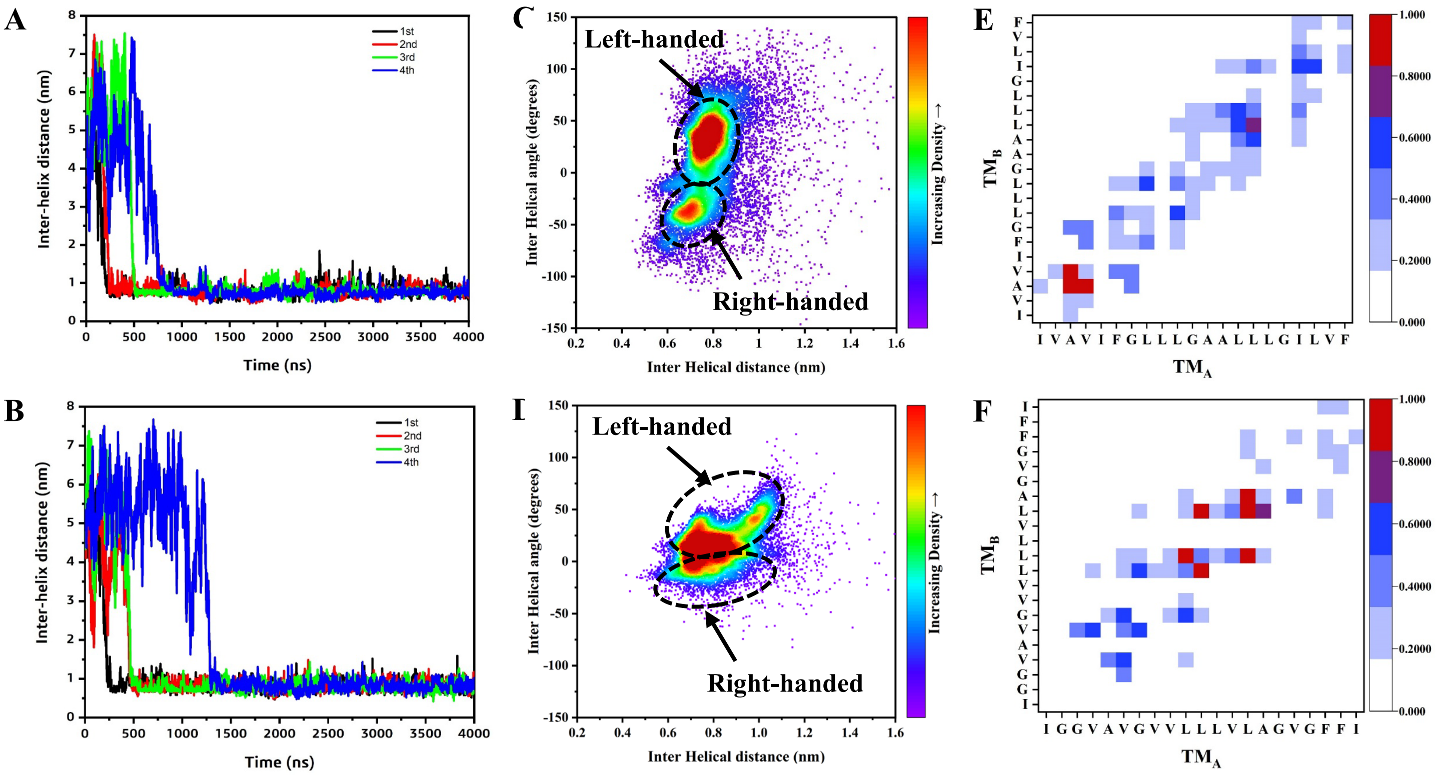


**Fig S2.** Comparison of the association of TM regions in the TM-only dimers of EphA1 and EphA2 in the mixed lipid bilayer. Left panel- Inter-helical distance plots showing the association between the TM regions of EphA1 (A) and EphA2 (B). 1st, 2nd, 3rd and 4th simulation results are shown black, red, green and blue lines. Middle panel- 2D distribution plot (interhelix angle vs. distance) for the EphA1 (C) and EphA2 (D). Data from the last 2.5μs simulations are considered for all the 4 simulations. Right panel- Contact map interface between the TMs for EphA1 (E) and EphA2 (F). Data from the last 2.5μs simulations are considered for all the 4 simulations. Contact maps are calculated with a cut off 5Å. The color scale (white to blue to red) indicates the fractional occupation of TM contacts (0 to 1). $TM_A$ and $TM_B$ denote the TM regions of chain A and chain B respectively.

**Figure S3.**

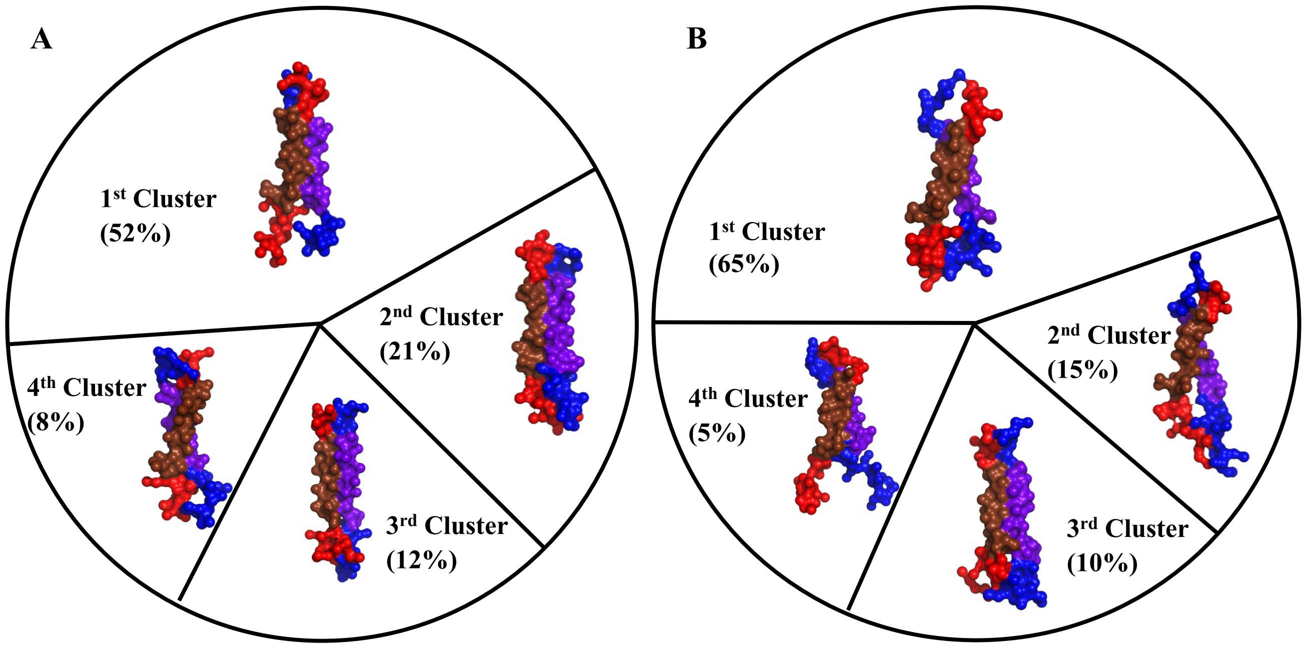


**Fig S3.** Comparison of most populated clusters of the TM-only dimer of EphA1 (A) and EphA2 (B) . All the conformations are shown in surface representation. The monomers are assigned as chain A and chain B throughout the paper. Chain A: N- & C-ter regions are colored as red and TM regions are colored as chocolate. Chain B: N- & C-ter regions are colored as blue and TM regions are colored as purple. Clustering of both EphA1 and EphA2 dimers are done with 6Å cutoff. Data from the last 2.5μs simulations are considered for all the 4 simulations.

### Figure S4.

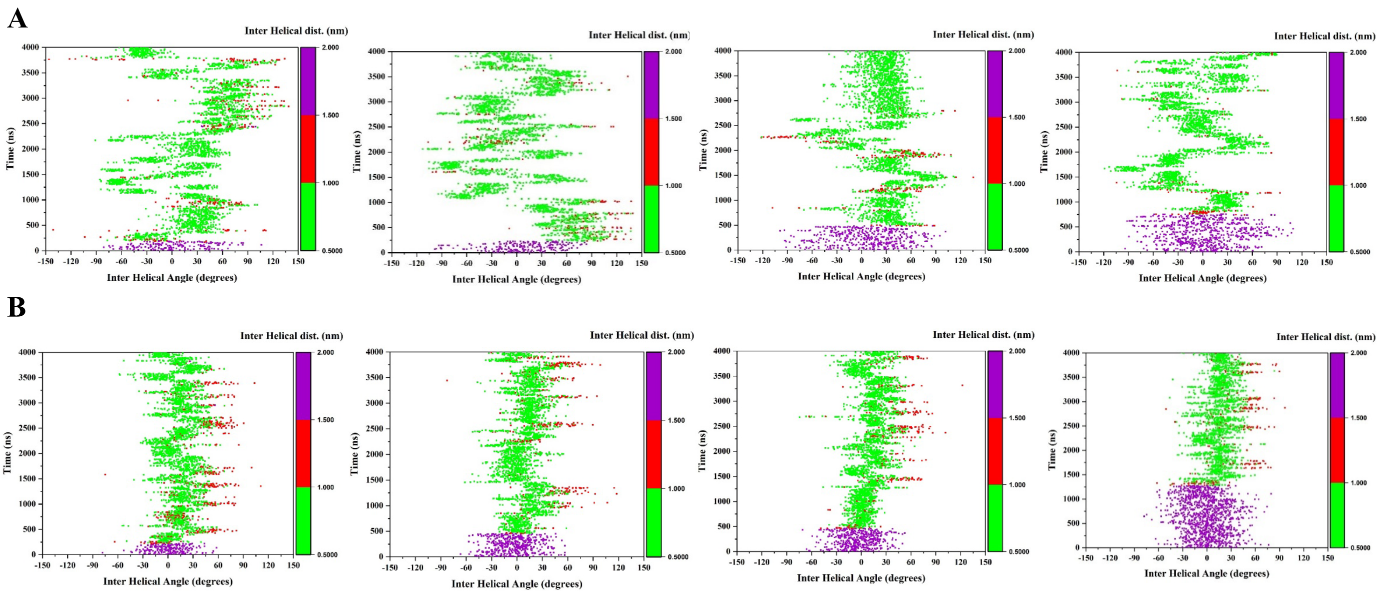


**Fig S4.** 2D Plots showing the conformational transition of TM-only dimers over the simulation time considering the inter-helical angle vs inter-helical distance for (A) EphA1 and (B) EphA2. Results from 4 trajectories are shown here. The plots are colored based on the inter-helical distance from 0.5- 1.0 nm (green), 1- 1.5 nm (red) and 1.5- 2.0 nm (purple).

#### Figure S5.

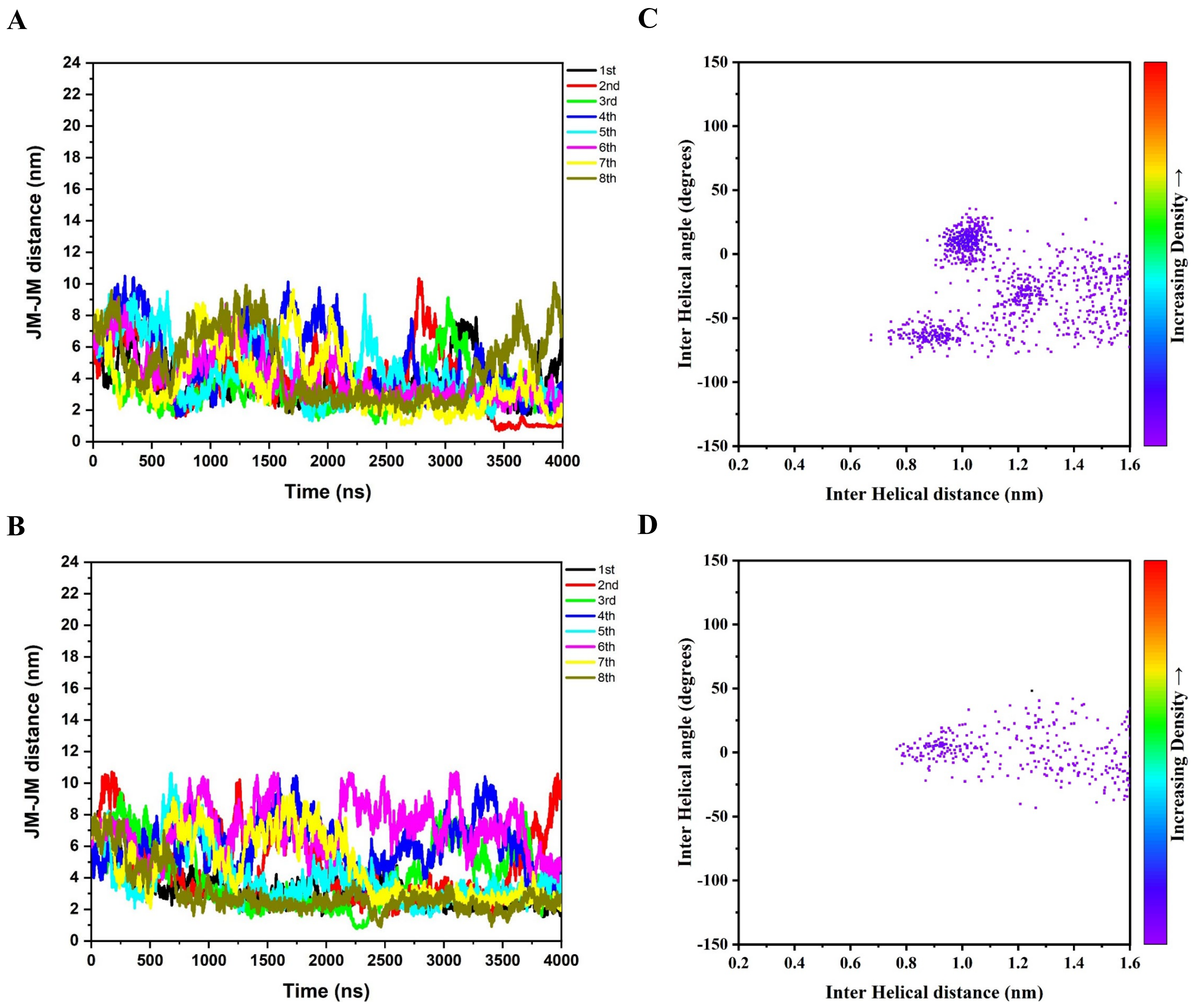


**Fig S5.** Comparison of the association of the TM regions in the FN2-TM-JM dimers of EphA1 and EphA2 in the mixed lipid bilayer. Left panel- Inter-helical distance plots showing the association between the TM regions of EphA1 (A) and EphA2 (B). $1^{st}$, $2^{nd}$, $3^{rd}$ and $4^{th}$ simulation results are shown black, red, green and blue lines. Right panel- 2D distribution plot (interhelix angle vs. distance) for the EphA1 (C) and EphA2 (D).

**Figure S6.**

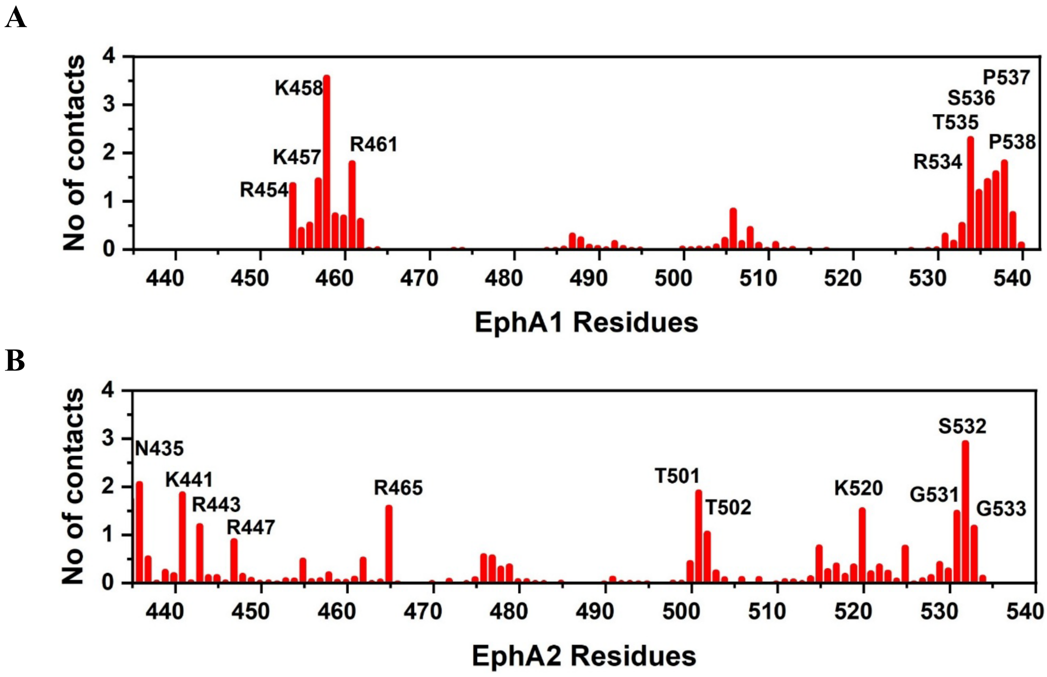


**Fig S6.** Comparison of the average PIP2 contacts with the FN2 domains of EphA1 (A) and EphA2 (B) for the FN2-TM-JM construct . Data from all eight replica simulations were used for the analysis. Contacts were defined using a 0.7 nm cutoff distance.

**Figure S7.**

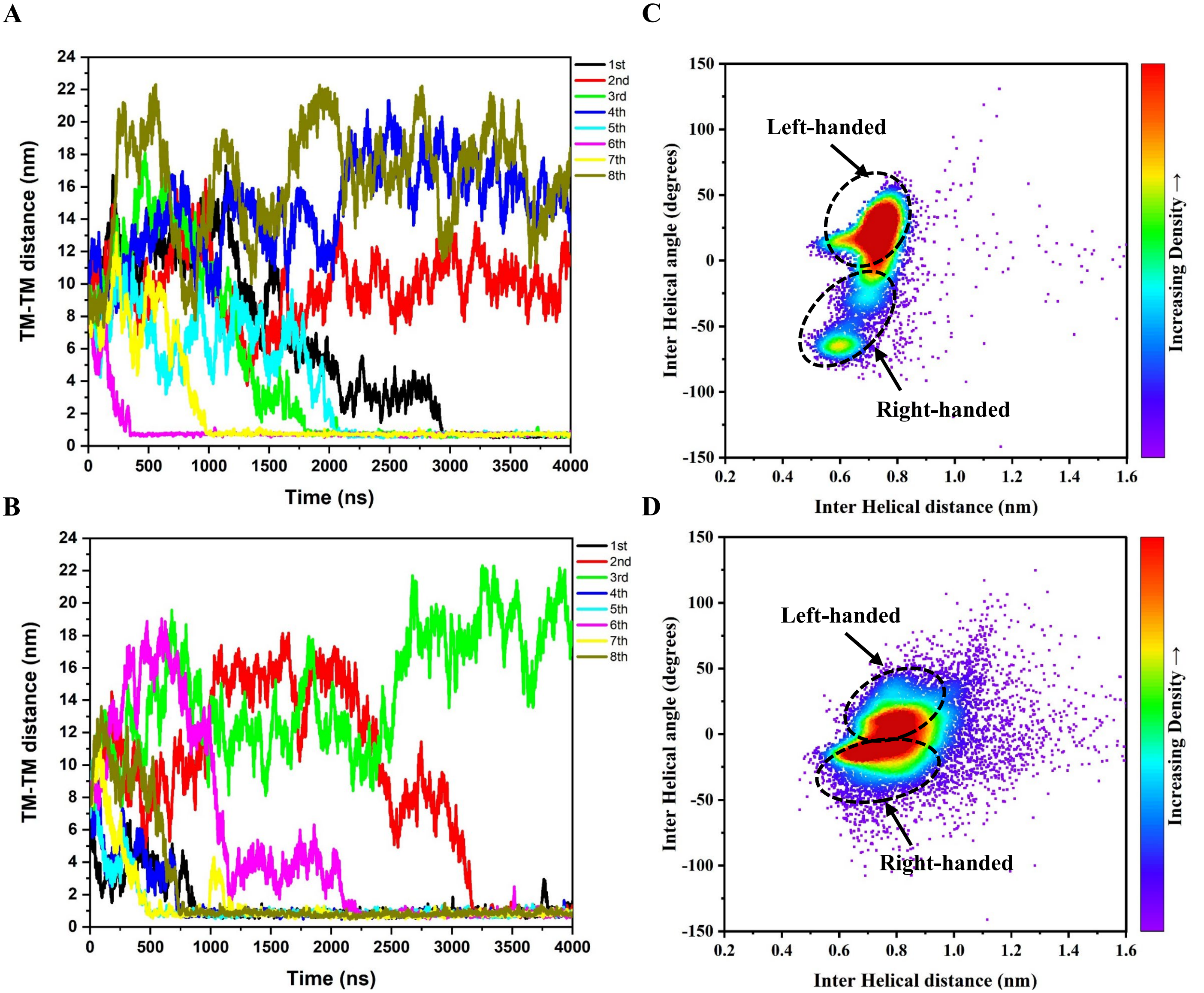


**Fig S7.** Comparison of the association of the TM regions in the FN1-FN2-TM-JM dimers of EphA1 and EphA2 in the mixed lipid bilayer. Left panel- Inter-helical distance plots showing the association between the TM regions of EphA1 (A) and EphA2 (B). Results from all the 8 simulations are shown in different colors. Right panel- 2D distribution plot (interhelix angle vs. distance) for the EphA1 (C) and EphA2 (D).

**Figure S8.**

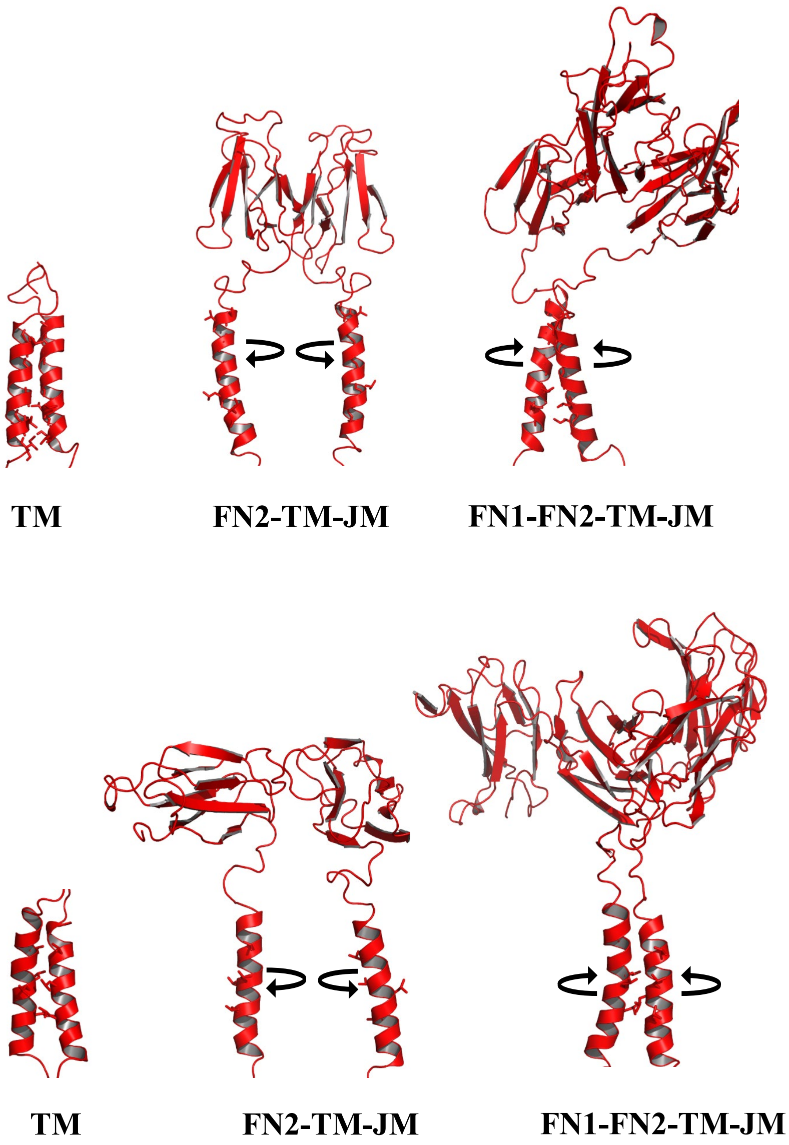


**Fig S8**. Comparison of major cluster of the all the three construct dimers of EphA1 (upper panel) and EphA2 (lower panel). All the conformations are shown in cartoon representation showing the TM interface residues. In case of the FN2-TM-JM construct, the interaction of FN2 domains leads to rotation of the interface residues resulting in no TM interaction compared to other constructs.

**Figure S9.**

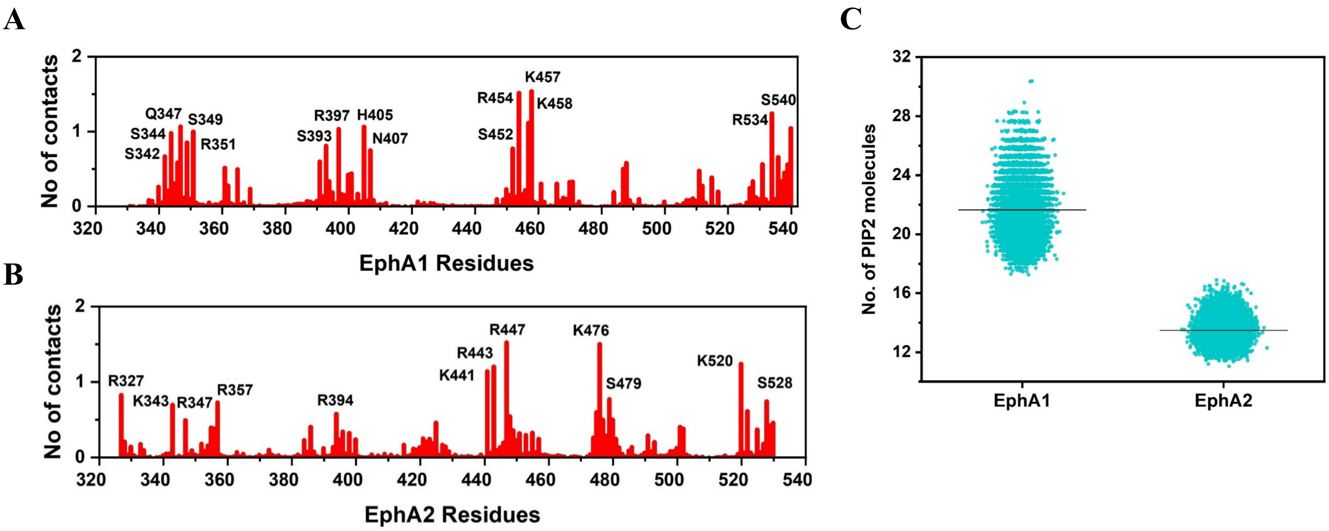


**Fig S9.** Comparison of the average PIP2 contacts with the FN domains of EphA1 (A) and EphA2 (B) for the FN1-FN2-TM-JM construct. Comparison of the PIP2 clustering around EphA1 and EphA2 (C). Data from all the 8 replica simulations are considered for calculation. We used 0.7 nm cutoff distance for contact calculation.

### Figure S10.

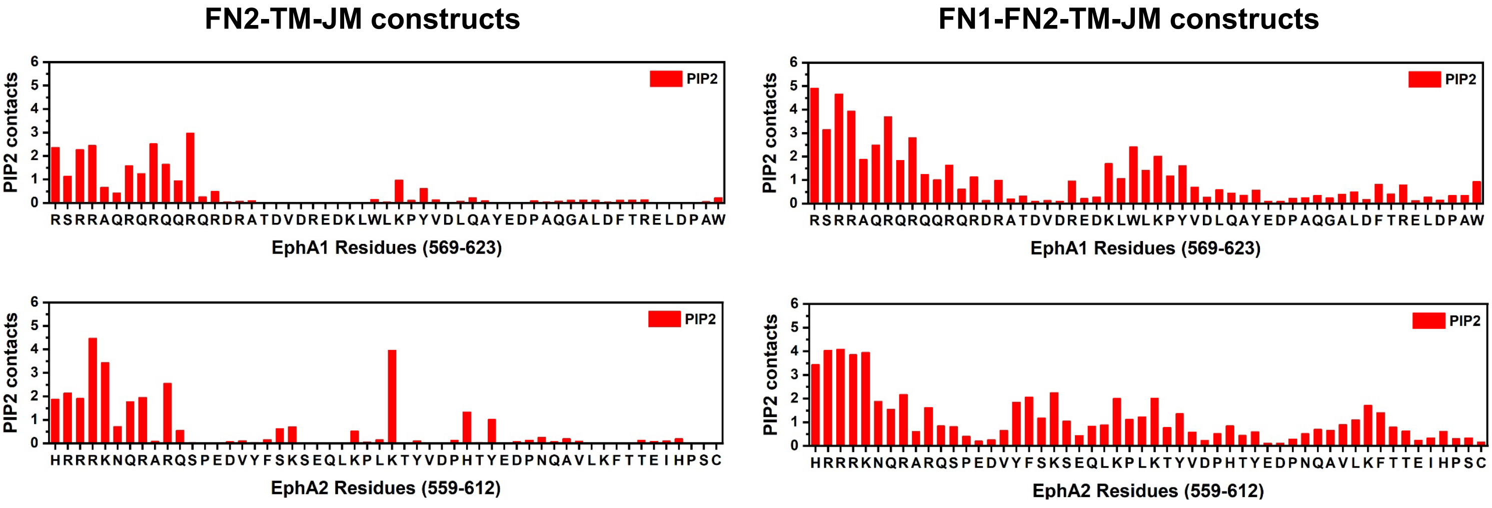


**Fig S10.** Comparison of PIP2 contacts with the JM regions for EphA1 and EphA2. Left panels show PIP2 contacts for the FN2-TM-JM dimers whereas right panels show contacts for the FN1-FN2-TM-JM constructs. Data from all the 8 replica simulations are considered for calculation. We used 0.7 nm cutoff distance for contact calculation.

### Figure S11.

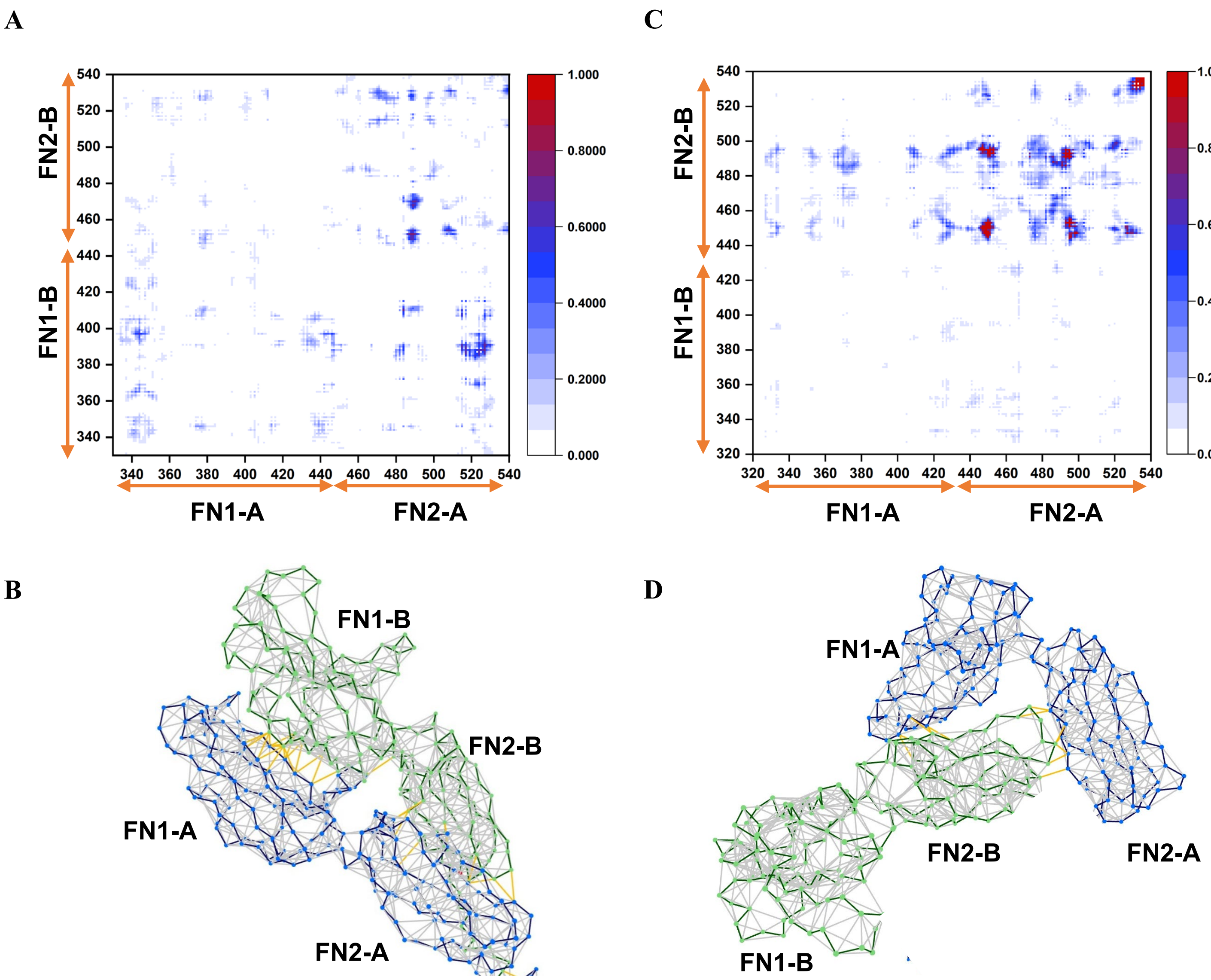


**Fig S11.** Comparison of contact interface between the FN1-FN2 domains in EphA1 (A) and EphA2 (C). Representations of the inter (shown in yellow lines) and intra- molecular (shown in grey lines) contacts in EphA1 (B) and EphA2 (D). We used 1.2 nm cutoff distance for contact calculation. Domains of chainA: FN1-A, FN2-A; domains of chain B: FN1-B, FN2-B.

**Figure S12.**

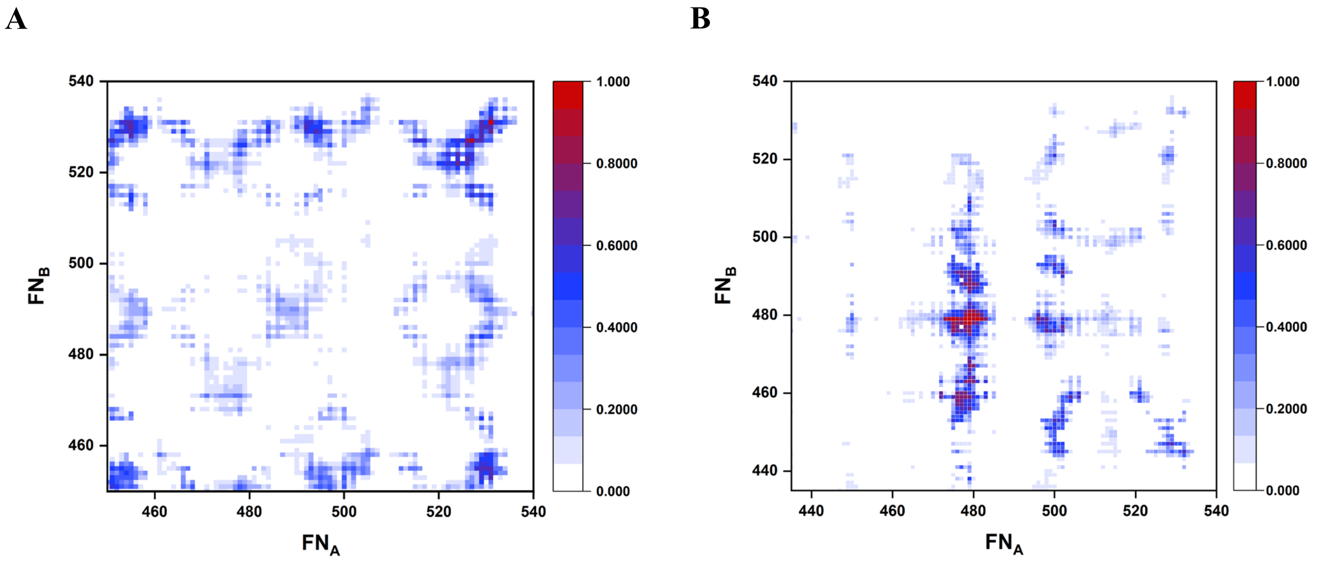


**Fig S12.** Comparison of contact interface between the FN2 domains in EphA1 (A) and EphA2 (B) for the FN2-TM-JM constructs.

**Figure S13.**

A

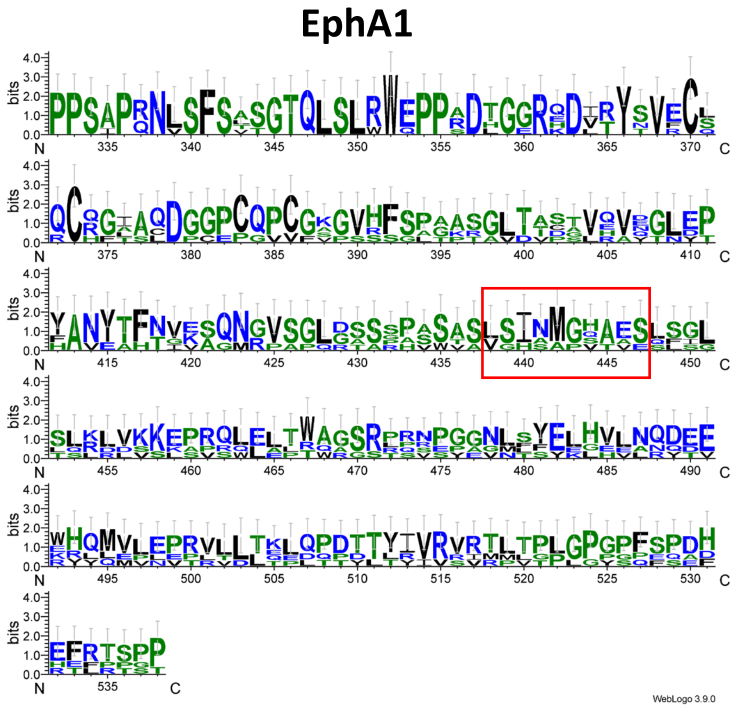


B

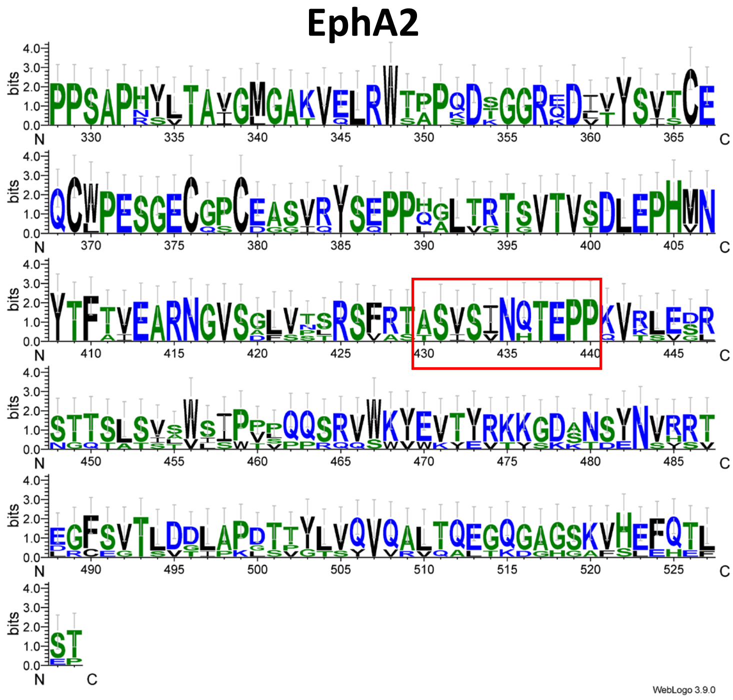


**Fig S13.** WebLogo representation of the multiple sequence alignment of the FN regions of EphA1 (A) and EphA2 (B). The y-axis indicates the information content (bit score), where a value of 4 corresponds to complete (100%) residue conservation at a given position. The x-axis denotes residue positions in the alignment. The linker regions are highlighted in the red box.

**Figure S14.**

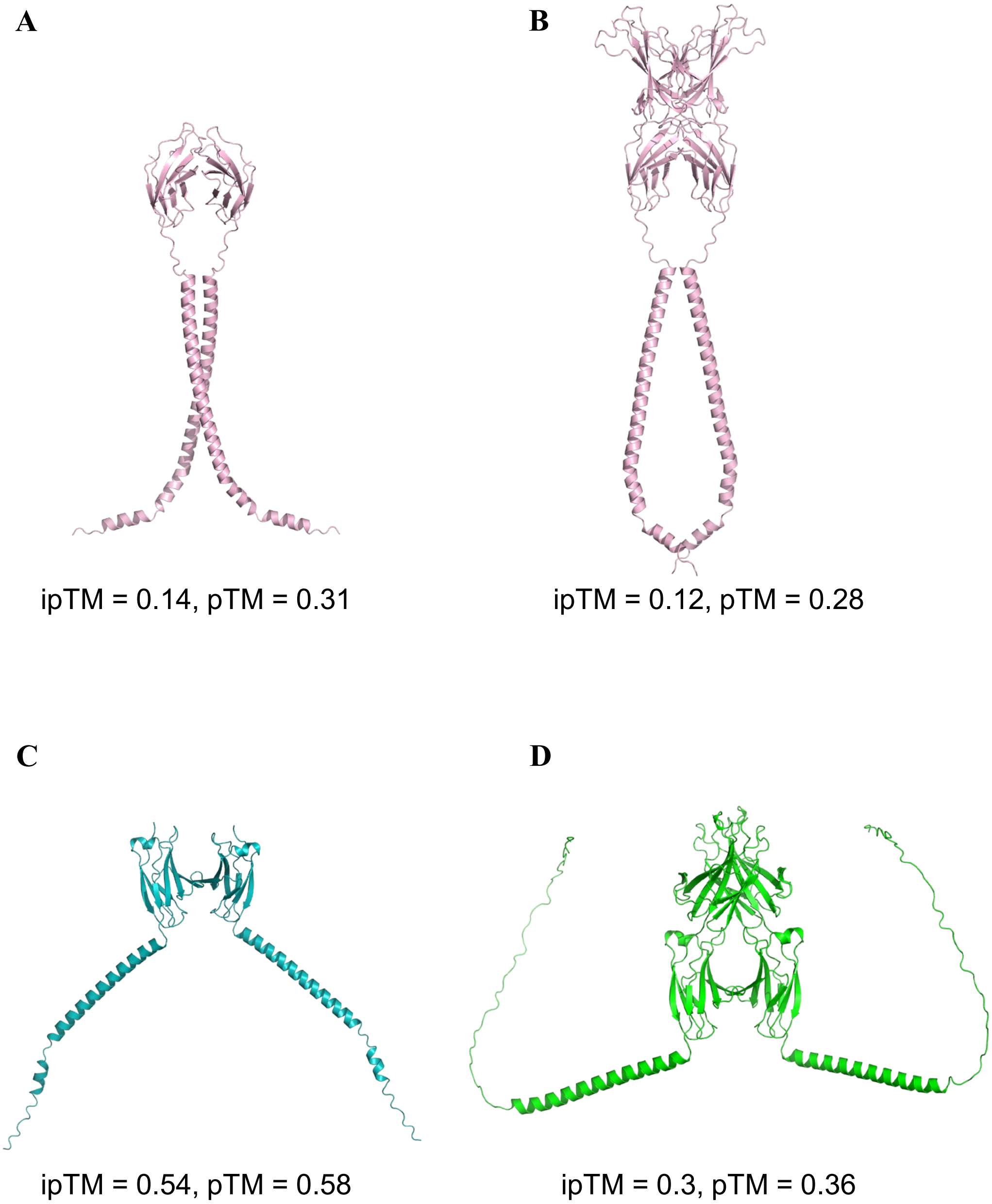


**Fig S14.** Alphafold3 predictions of the homodimers of FN2-TM-JM and FN1-FN2-TM-JM constructs for EphA1 (A-B) and EphA2 (C-D).
